# Neurotrophin receptor activation rescues cognitive and synaptic abnormalities caused by mutation of the psychiatric risk gene *Cacna1c*

**DOI:** 10.1101/2020.05.29.123695

**Authors:** Cezar M. Tigaret, Tzu-Ching E. Lin, Edward Morrell, Lucy Sykes, Michael C. O’Donovan, Michael J. Owen, Lawrence S. Wilkinson, Matthew W. Jones, Kerrie L. Thomas, Jeremy Hall

## Abstract

Genetic variation in *CACNA1C*, which encodes the alpha-1 subunit of Ca_V_1.2 L-type voltage-gated calcium channels, is strongly linked to risk for psychiatric disorders including schizophrenia and bipolar disorder. To translate genetics to neurobiological mechanisms and rational therapeutic targets, we investigated the impact of altered *Cacna1c* dosage on rat cognitive, synaptic and circuit phenotypes implicated by patient studies. We show that rats hemizygous for *Cacna1c* harbor marked impairments in learning to disregard non-salient stimuli, a behavioral change previously associated with psychosis. This behavioral deficit is accompanied by dys-coordinated network oscillations during learning, pathway-selective disruption of hippocampal synaptic plasticity, attenuated Ca^2+^ signaling in dendritic spines and decreased signaling through the Extracellular-signal Regulated Kinase (ERK) pathway. Activation of the ERK pathway by a small molecule agonist of TrkB/TrkC neurotrophin receptors rescued both behavioral and synaptic plasticity deficits in *Cacna1c^+/-^* rats. These results map a route through which genetic variation in *CACNA1C* can disrupt experience-dependent synaptic signaling and circuit activity, culminating in cognitive alterations associated with psychiatric disorders. Our findings highlight targeted activation of neurotrophin signaling pathways with BDNF mimetic drugs as a novel, genetically informed therapeutic approach for rescuing behavioral abnormalities in psychiatric disorder.

**One Sentence Summary:** Neurotrophin receptor activation reveals that BDNF mimetic drugs have therapeutic potential to ameliorate genetic risk for psychiatric disorders.

## Introduction

The major psychiatric disorders such as schizophrenia and bipolar disorder place an enormous burden on society. These conditions have not however seen the advances in mechanistic understanding and therapy realized in other areas of medicine. There is now hope that recent advances in the understanding of the genomic basis of these conditions may pave the way to the development of new therapeutics (*1*). Particularly promising in this respect is the demonstration of a strong association between genetic variation in voltage gated calcium channels with schizophrenia and bipolar disorder (*2*). Genome-wide association studies (GWAS) have consistently identified single nucleotide polymorphisms (SNPs) in *CACNA1C*, which encodes the pore-forming α_1C_ subunit of Ca_V_1.2 L-type VGCCs (L-VGCCs), as having significant association with both conditions (*3, 4*). While schizophrenia and bipolar disorder can present differently in the clinic, both are associated with psychosis, and genomic studies have indicated a significant shared genetic architecture between the two disorders (*5*).

The exact molecular effects of the *CACNA1C* SNPs implicated by GWAS are not yet fully understood. However, risk-associated common variants in *CACNA1C* are intronic and are likely to act by altering gene expression. Previous studies have indicated that these risk variants may act to decrease expression of *CACNA1C* (*6, 7*), including in the human hippocampus (*8*). The association of rare deleterious mutations in genes encoding VGCC subunits, including *CACNA1C*, in people with schizophrenia and other neurodevelopmental disorders, further supports the view that decreased expression of *CACNA1C* can contribute to disease risk (*9, 10*). Therefore, understanding the effects of reduced *CACNA1C* dosage is important in discerning how genetic variation in L-VGCCs might contribute to risk for neuropsychiatric illness.

Ca_V_1.2 L-VGCCs are highly expressed the mammalian brain, including in the hippocampus (*11*). Clinical and preclinical studies have implicated hippocampal structural and functional abnormalities in the pathophysiology of schizophrenia and bipolar disorder (*12–14*). Impairments of hippocampal-dependent associative learning processes have also been considered to contribute to the development of psychosis (*15–21*).

Associative learning in the hippocampus is orchestrated by rhythmic neural activity in the entorhinal-hippocampal network and is underpinned by associative synaptic plasticity at hippocampal glutamatergic synapses (*22–24*). Induction of plasticity at these synapses relies on the coordinated activation of both postsynaptic NMDA receptors (NMDAR) and voltage-gated calcium channels including L-VGCCs (*25–27*).

Neuronal L-VGCCs play a key role in linking membrane depolarization to the activation of signaling pathways in particular the Extracellular-signal Regulated Kinase (ERK) signaling pathway. The ERK pathway critically regulates both early (*28, 29*) and late stages of synaptic plasticity (*30, 31*). L-type VGCC-dependent ERK signaling activates transcription factors including CREB that control the expression of genes required for long-lasting plasticity, including the neurotrophin BDNF (Brain Derived Neurotropic Factor) (*30, 31*). Thus, calcium entry through L-VGCCs plays a critical role in regulating the changes in synaptic efficacy and gene expression underlying associative learning.

In the present study, we used a translationally relevant rat model, *Cacna1c^+/-^* to examine the impact of reduced dosage of the psychiatric risk gene *Cacna1c* on hippocampus-dependent associative learning and went on to reveal the impact of reduced *Cacna1c* dosage on underlying synaptic plasticity, network synchronization and ERK pathway signaling in the hippocampus. Having mapped the links between genetic variation in *CACNA1C* and disruptions to experience-dependent synaptic signaling, circuit activity and behavior we then showed that activation of the ERK signaling pathway through a small molecule agonist of the TrkB/TrkC neurotrophin receptors could rescue impairments in both behavior and synaptic plasticity. Our findings suggest that targeted activation of neurotrophin signaling pathways with BDNF mimetic drugs may be of use in treating cognitive and brain abnormalities in psychiatric disorder.

## Results

### Behavioral effects of Cacna1c hemizygosity on latent inhibition

We investigated the effects of reduced Ca_V_1.2 dosage on associative learning using rats hemizygous for a truncating mutation in exon 6 of *Cacna1c* gene encoding the pore-forming α_1C_ subunit of Ca_V_1.2 L-VGCCs (*32*) (Methods). *Cacna1c*^+/-^ rats have an approximately 50% decrease in both *Cacna1c* mRNA and protein in the hippocampus, mimicking the anticipated impact of deleterious mutations in *CACNA1C* in humans (*33*). We assessed the behavior of *Cacna1c^+/-^* rats and wild type littermates using tests of contextual fear conditioning (CFC), as the formation of contextual fear associations is known to depend on the hippocampus (*34*) and to require L-VGCCs (*35*). To determine the ability of *Cacna1c^+/-^* rats to establish and retrieve contextual fear associations we used a paradigm where animals received a single mild footshock (US, 2s, 0.5 mA) 2 min after being placed in a novel context (Fig. 1A and Supplementary Methods). Memory for the context-fear association was assessed by measuring the freezing response upon return to the conditioned context 3h (short-term memory, STM), 24h (long-term memory, LTM1), and 8 days (LTM2) after CFC training. *Cacna1c^+/+^* and *Cacna1c^+/-^* animals had equivalent levels of fear response at all recall sessions, as shown in Fig. 1A and replicated in a separate cohort (Fig. S1), indicating that reduced dosage of *Cacna1c* has no impact on contextual fear learning *per se*.

**Fig. 1.**
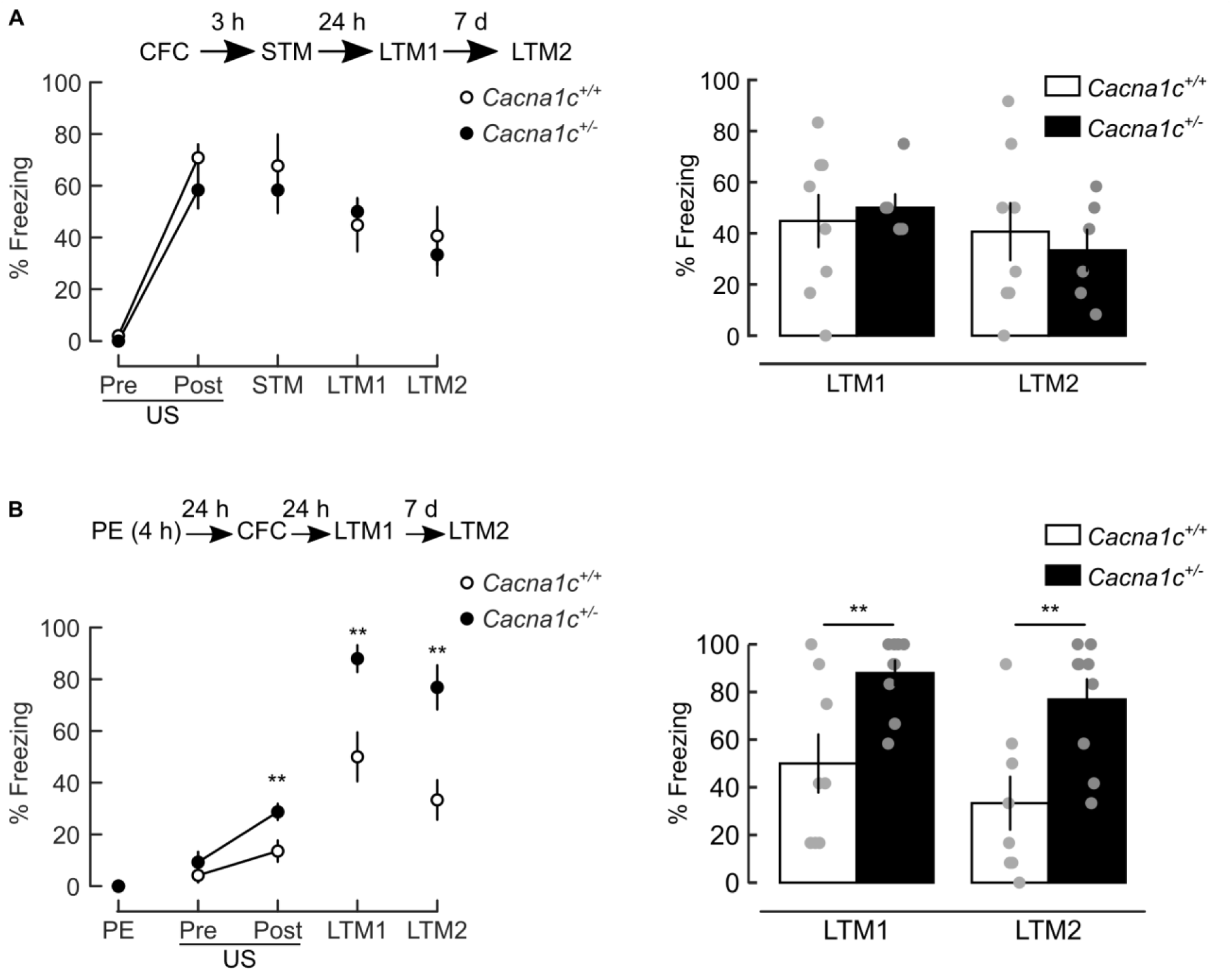
Low *Cacna1c* dosage selectively impairs latent inhibition (LI) of contextual fear memory CFM. **All panels**: *Left*: schematics of behavioral protocols (*top*) and freezing responses measured during all trials (*bottom*). PE: 4h context pre-exposure; CFC: contextual fear conditioning comprising Pre-US and Post-US epochs before and after footshock (US), respectively; STM, LTM1, LTM2: short-term and long-term contextual fear memory (CFM) retrieval trials. *Right*: Summary of freezing responses at LTM tests. (**A**) Acquisition and consolidation of CFC are unaltered in *Cacna1c^+/-^* animals. Effects of genotype: F_(1,12)_=0.385, p=0.546, trial: F_(4,48)_=29.624, p<0.001; genotype × trial interaction: F_(4,48)_=0.533, p=0.712; *Cacna1c^+/-^*: n=6; *Cacna1c^+/+^*: n=8). (**B**) Higher levels of freezing post US and during CFM retrieval trials indicate the *Cacna1c^+/-^* animals have a deficit in the LI. Genotype × session interaction: F_(2.1,31.49)_=6.119, p=0.005; pairwise comparisons between genotypes: Post-US: F_(2.1,31.47)_=10.464, p=0.006; LTM1: F_(2.1,31.49)_=8.895, p=0.009; LTM2: F_(2.1,31.49)_=9.869, p=0.007(Greenhouse–Geisser corrected); *Cacna1c^+/-^*: n=9; *Cacna1c^+/+^*: n=8. Data presented as means ±SEM. **p <0.01 determined by two-way repeated measures ANOVA followed by pairwise post-hoc comparison with Bonferroni correction.

We next assessed the performance of *Cacna1c^+/-^* rats and wild-type littermates in a paradigm of latent inhibition (LI) of contextual fear conditioning (schematics in Fig. 1B and Fig. S1). LI is the reduced ability to form conditioned associations to a stimulus (in this case, the conditioning context) as a result of pre-exposure to the stimulus alone and reflects the normal process of learning to ignore irrelevant stimuli. Notably latent inhibition has previously been found to be impaired in individuals experiencing psychosis (*15, 16*). For LI training, animals were preexposed to the to-be-conditioned context (PE) for 4h, then subjected to CFC training 24h or 48h later (Figs. 1B, S1). In *Cacna1c^+/+^* animals, the preexposure resulted in a robust LI of CFC manifested as reduced freezing response at long-term memory sessions (Figs. 1B, S1). In contrast, *Cacna1c^+/-^* animals had a marked deficit in LI of CFC, an effect seen in two separate cohorts (Fig. 1B, S1). These results show that *Cacna1c* hemizygosity selectively impairs the LI of contextual fear associations. To determine whether this effect was mediated through an impact on L-VGCCs specifically in the hippocampus we infused the L-VGCC antagonist diltiazem (DTZ) into the dorsal hippocampus during context pre-exposure and observed a similar impairment in the establishment of latent inhibition (Fig. S2). Taken together, these results indicate a specific role of hippocampal Ca_V_1.2 channels during the preexposure stage in the LI of CFC paradigm.

LI of CFC requires the ability of dorsal hippocampus to form and store context-specific memories (*36*). We hypothesized that the deficit in LI of CFC observed in *Cacna1c^+/-^* animals reflects a disruption of dorsal hippocampal processes that form representations of a novel environment during preexposure. Therefore we investigated the impact of *Cacna1c* hemizygosity on two fundamental neural mechanisms proposed to support memory encoding in the hippocampus: associative plasticity at CA1 pyramidal cell synapses (*24*) and phase-amplitude coupling between the theta and gamma oscillations of the local field potential in CA1 (*22*).

### Cacna1c^+/-^ rats show altered plasticity in the dorsal hippocampal CA1

Formation of stable context-specific representations in the hippocampus requires strengthening of cortical excitatory inputs to CA1 pyramidal neurons. These inputs arrive indirectly through the CA3 subfield via the Schaffer collateral (SC-CA1) pathway, and directly through the temporo-ammonic (TA-CA1) pathway (*37–40*). We examined the induction of LTP at SC-CA1 and TA-CA1 synapses onto CA1 pyramidal neurons in the dorsal hippocampus using *ex vivo* slices from *Cacna1c*^+/-^ and *Cacna1c*^+/+^ rats. To induce LTP we used a theta-burst pairing protocol consisting of synaptic stimulation coincident with post-synaptic action potentials (TBP, Fig. 2A). TBP mimics neuronal activity patterns observed during learning *in vivo* (*41*) and relies on the coordinated activation of postsynaptic NMDAR and VGCCs (*27, 42*). TBP failed to induce LTP at SC-CA1 synapses in slices from *Cacna1c*^+/-^ animals (Fig. 2B,D) whereas it produced robust SC-CA1 LTP in *Cacna1c*^+/+^ slices (Fig. 2C,D). In *Cacna1c*^+/+^ slices the TBP-induced LTP at SC-CA1 synapses was also blocked by L-VGCC antagonists isradipine, DTZ, and the NMDAR glycine site antagonist L-689560 (Fig. S3A-C,H), confirming that TBP-induced LTP requires the activation of both NMDARs and L-VGCCs.

**Fig. 2.**
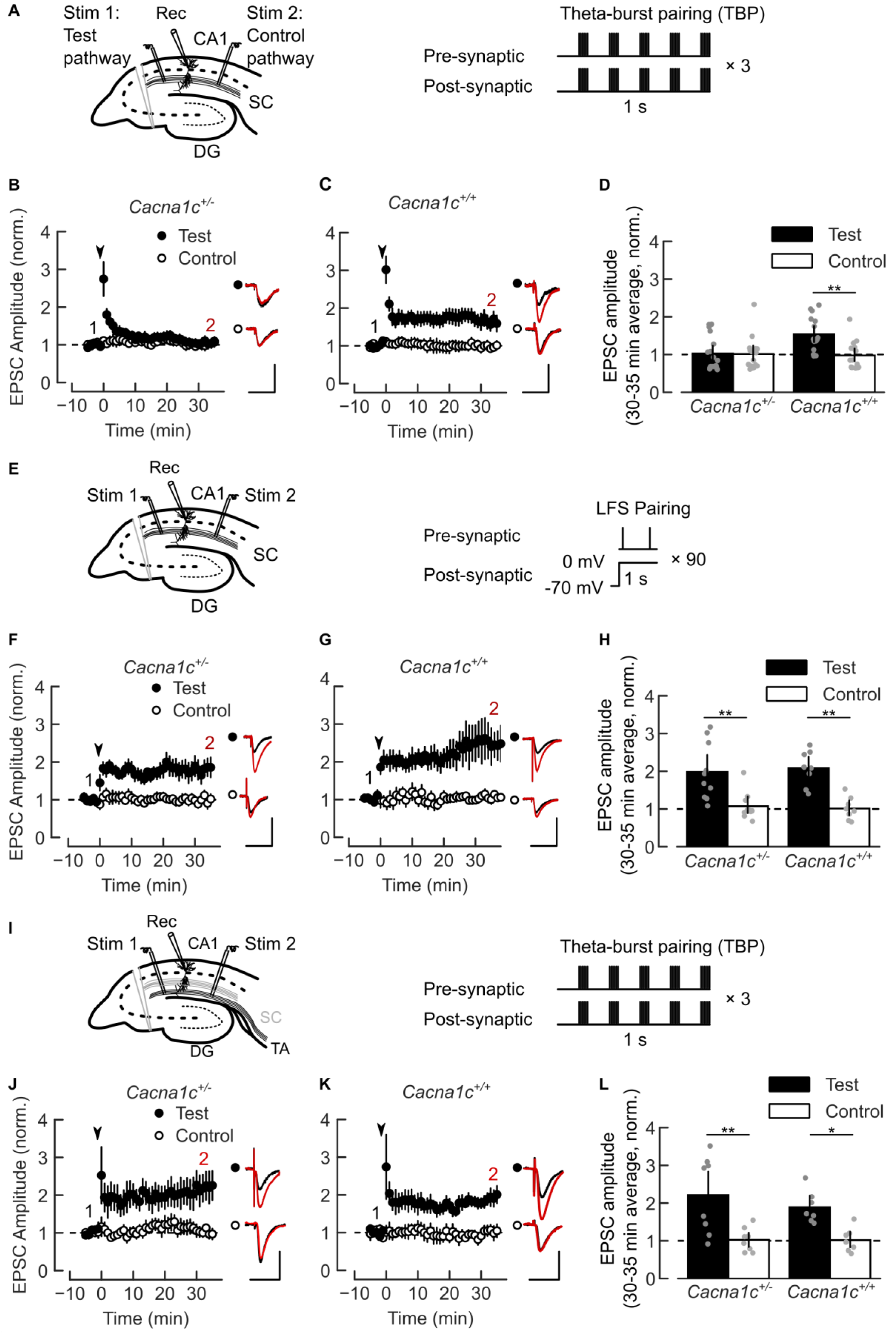
Reduced *Cacna1c* dosage impairs LTCC-sensitive induction of associative LTP selectively at Schaffer collateral to CA1 (SC-CA1) synapses. (**A**) Electrodes placements (*left*) and theta-burst pairing (TBP) protocol (*right*) for LTP at SC-CA1 synapses. (**B**) No TBP-induced LTP at SC-CA1 synapses in *Cacna1c*^+/-^slices. (**C**) Robust TBP-induced homosynaptic LTP at SC-CA1 in *Cacna1c*^+/+^ slices. (**D**) Summary of normalized change in EPSC amplitude at 30-35 min shown in B and C. Effect of genotype on LTP: LR_(1)_=5.16, p=0.023; genotype × pathway interaction: LR_(1)_=7, p=0.0081). Pairwise comparisons of Test vs Control pathway for *Cacna1c^+/-^*: p=0.99, n=17/12; *Cacna1c^+/+^*: p=0.0057, n=16**/**9. (**E**) Electrodes placements (*left*) and low-frequency stimulation (LFS-pairing) protocol (*right*) for SC-CA1 LTP. (**F,G**) LFS-pairing induces SC-CA1 LTP in *Cacna1c^+/-^* (F) and *Cacna1c*^+/+^ slices (G). (**H**) Summary of EPSC amplitude change at 30-35 min shown in F and G: no effect of genotype (LR_(1)_=0.103, p=0.748) or genotype × pathway interaction (LR_(1)_=0.227, p=0.633) on LTP. Pairwise Test vs Control for *Cacna1c*^+/-^: p=0.0027, n=10/7; *Cacna1c*^+/+^: p=0.0011, n=8/6. (**I**) Electrode placements (*left*) and TBP protocol (*right*) for LTP at TA-CA1 synapses. (**J,K**) TBP induces homosynaptic TA-CA1 LTP in slices from *Cacna1c^+/-^* (J) and *Cacna1c^+/+^* animals (K). (**L**) Summary of changes in normalized EPSC amplitude at 30-35 min shown in F and G: no effect of genotype (LR_(1)_=0.0258, p=0.8724) and no genotype × pathway interaction (LR_(1)_=0.065, p=0.798); contrast Test *versus* Control pathways, in *Cacna1c^+/-^*: p=0.0093, n=8/7; *Cacna1c^+/+^*: p=0.014, n=7/5. Plots show EPSC amplitude time-course in Test and Control pathways, normalized to 5 min average before LTP induction (arrowheads). Insets: 5 min average EPSC waveforms before (1, black) and 30-35 min after (2, red) LTP induction; scale bars: 50 pA, 50 ms. Sample sizes given as cells/animals. Data presented as means ±SEM. *p <0.05, **p<0.01 determined by two-way ordinal regression (cumulative link model) followed by analysis of deviance (ANODE).

The deficit of TBP-induced LTP at SC-CA1 synapses in *Cacna1c*^+/-^ animals could be selective or could reflect a broad impairment of plasticity mechanisms. To disambiguate these possibilities, we tested the induction of SC-CA1 LTP by an alternative protocol of low frequency pre-synaptic stimulation paired with tonic post-synaptic depolarization (LFS-pairing, Fig. 2E). LFS-pairing induces NMDAR-dependent synaptic potentiation with no contribution from post-synaptic action potentials or L-VGCCs (*43*). We observed a robust SC-CA1 LTP induced with LFS-pairing in both genotypes (Fig. 2F-H). In *Cacna1c*^+/+^ slices LFS-pairing-induced LTP was insensitive to either isradipine or DTZ but was blocked by L-689560 (Fig. S3D-F,H) confirming that NMDAR activation but not L-VGCC activation is required for this form of LTP. In contrast to SC-CA1 synapses, TBP induced a robust LTP at TA-CA1 synapses in both genotypes (Fig. 2I-L) which was sensitive to L-VGCC block in wild-type slices (Fig. S3G,H).

Overall, these results show that *Cacna1c* hemizygosity impairs forms of LTP that depend on the activation of post-synaptic L-VGCCs during somatic action potential bursts without affecting NMDAR-dependent mechanisms of synaptic plasticity, at the SC-CA1 pathway.

### CA1 neurons in Cacna1c^+/-^ rats have impaired spine Ca^2+^ signaling associated with postsynaptic spike bursts

LTP requires a neuronal associative signal which, in hippocampal neurons, can be provided by postsynaptic action potentials (AP) backpropagated at SC-CA1 synapses and by local dendritic spikes at the more distal TA-CA1 synapses (*26, 44*). Therefore, we investigated the excitability of CA1 pyramidal neurons in hippocampal slices from *Cacna1c*^+/-^ and *Cacna1c*^+/+^ animals. The passive membrane properties, AP threshold and rheobase current were comparable in CA1 pyramidal neurons from both genotypes (Supplementary Table S1, Fig. S4A-E). However, *Cacna1c*^+/-^ neurons showed a significant reduction in the broadening of somatic AP waveforms during high frequency (>40Hz) AP firing when compared to wild-type cells (Fig. S4F-I). In *Cacna1c*^+/+^ neurons the L-VGCC blocker isradipine inhibited somatic AP broadening to levels comparable to those in *Cacna1c*^+/-^ neurons in the absence of the drug (Fig. S4J,K). Somatic AP broadening occurs normally during high-frequency AP bursts and reflects the slowing of AP repolarization mediated by voltage- and Ca^2+^-sensitive K^+^ channels (*45, 46*). The impaired broadening of somatic APs in *Cacan1c^+/-^* pyramidal neurons indicates that low Ca_V_1.2 dosage alters the Ca^2+^-sensitive spike repolarization during burst firing.

Somatic AP broadening in CA1 pyramidal neurons has been proposed to facilitate the dendritic backpropagation of somatic spikes (*47*) necessary for the induction of associative synaptic plasticity such as TBP-LTP (*27, 42, 44*) and to increase the gain of intracellular Ca^2+^ signals associated with AP bursts (*45*) such as those occurring during the exploration of a novel environment (*40, 48*). The reduced AP broadening during burst firing in *Cacna1c*^+/-^ neurons may impact on postsynaptic Ca^2+^ signals triggered by APs backpropagated in dendritic spines during plasticity induction. We tested this hypothesis by comparing Ca^2+^ transients elicited with high-frequency somatic AP bursts in spines located on radial oblique dendrites, in *Cacna1c*^+/-^ vs *Cacna1c*^+/+^ CA1 pyramidal neurons (Fig. 3A,B, S5, Supplementary Tables S2, S3).

**Fig. 3.**
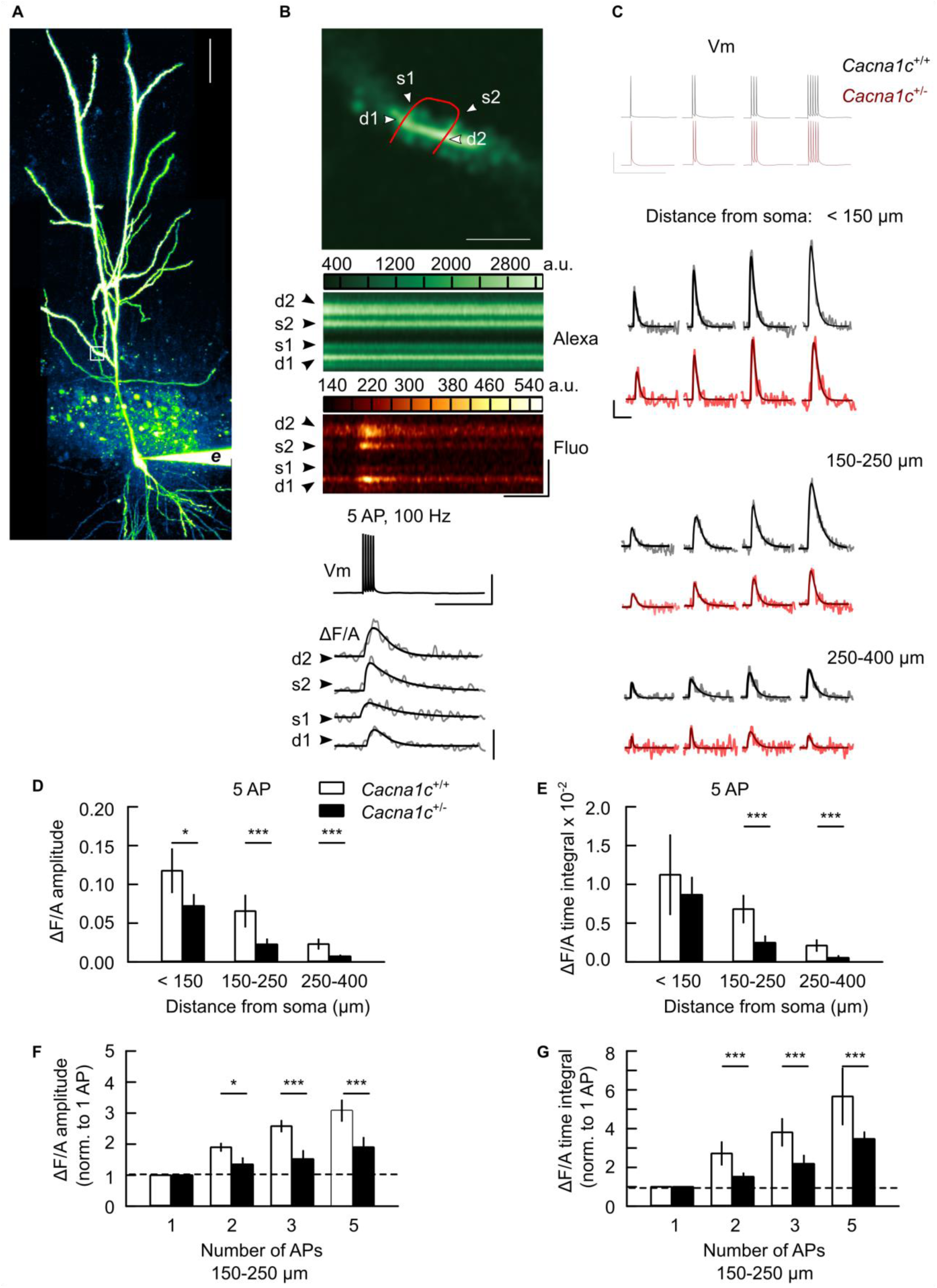
Low *Cacna1c* dosage is correlated with a reduction of spine Ca^2+^ signals elicited by back-propagated action potentials in CA1 pyramidal cells. (**A**) CA1 pyramidal neuron during spine Ca^2+^ imaging (Z-projected pseudo-color image, Alexa Fluor 594 channel; white square: spine imaging region; *e*: recording electrode; scale: 50 µm). (**B**) Imaging of spine Ca^2+^ transients elicited by back-propagated action potentials (APCaTs). Top-to-bottom: imaging region in (A) with spines (s1, s2), parent dendrite (d1, d2) and line-scan trajectory (red); line-scan images, Alexa Fluor 594 (Alexa) and Fluo 5F (Fluo) channels, during five somatic APs (Vm); APCaTs during the 5 APs (grey: ΔF/A, change in fluorescence intensity in Fluo channel, relative to Alexa channel; black: double-exponential fit). Scales, vertical: 2 µm, 20 mV, 0.1 arbitrary units (a.u.); horizontal: 0.2 s. (**C**) Example APCAT waveforms elicited by 1-5 somatic AP (Vm) in spines within three distance zones. Scale: 20 mV, 0.02 a.u., 0.2 s. (**D,E**) APCaTs have smaller amplitude (D) and time integral (E) in *Cacna1c^+/-^ versus Cacna1c^+/+^* neurons (pairwise comparisons of 5 AP APCaTs, in distance zones (µm) < 150: amplitude p=0.0473, time integral p=0.414; 150-250: both measures p<0.0001; 250-400: both measures p<0.0001). (**F,G**) Summation of APCaT amplitude (F) and time integral (G) with the number of APs is reduced in *Cacna1c^+/-^ versus Cacna1c^+/+^* neurons (pairwise comparisons for APCaTs at 150-250 µm, for 2 AP: normalized amplitude p=0.15, normalized time integral p<0.0001, 3 AP: both measures p<0.0001; 5 AP: both measures p<0.0001). Data was expressed as means ± SEM. *p <0.05, **p < 0.01 and ***p< 0.001. determined by two-way repeated measures ANOVA followed by Tukey adjustment of p-values for contrasts. Statistical analysis results are given in Supplementary Table S2; all p-values for pairwise comparisons are given in Supplementary Table S3.

Backpropagated APs activate spine VGCCs including the L-type (*27, 49*). Spine Ca^2+^ transients elicited by back-propagated APs (APCaTs) attenuated with increasing distance from the soma (Fig. 3C-E, S6). However, in *Cacna1c*^+/-^ neurons the APCaTs in spines at 150-250 µm from the soma were consistently smaller compared to those in wild-type cells (Fig. 3D,E) and summated poorly with the number of APs per burst (Fig. 3F,G, S7).

These results show that the low Ca_V_1.2 dosage in *Cacna1c*^+/-^ CA1 pyramidal neurons alters the burst-associated spine Ca^2+^ signaling at SC-CA1 synapses, which may underlie the selective deficit in synaptic plasticity induction during burst firing and impaired TBP-LTP at SC-CA1 synapses.

### Cacna1c^+/-^ rats have reduced phase-amplitude coupling between theta and gamma oscillations of dorsal CA1 local field potential

During the encoding of novel environments, the induction of synaptic plasticity leading to the establishment of neuronal ensembles depends on the critical timing of excitatory inputs, coordinated by the synchronization of the rhythmic activity in the CA1 subfield (*22, 50, 51*). To determine the impact of *Cacna1c* heterozygosity on the synchronization of neural activity in the CA1 network, we monitored the modulation of the local field potential (LFP) gamma oscillations by the phase of LFP theta oscillations (theta-gamma Phase-Amplitude Coupling (PAC)) in the dorsal CA1. In the hippocampus, theta-gamma PAC has been proposed as a mechanism for the storage and recall of object and event representations (*23, 52, 53*). In the dorsal CA1, the slow (∼25-40 Hz) and fast (∼ 65-140 Hz) gamma sub-bands are thought to reflect the synchronization of CA1 pyramidal neurons with the activity of CA3 neurons via SC-CA1 pathway, and that of the medial entorhinal cortex (MEC) neurons, via TA-CA1 pathway, respectively (*54*).

To determine the changes in theta-gamma PAC associated with the exploration of a novel environment, we recorded LFP oscillations in dorsal CA1 in rats running along a track in a familiar, then a novel environment (Figs. 4A and S8A-C). In the familiar environment, animals from both genotypes had similar phase-amplitude coupling across the gamma frequency spectrum (Fig. 4B,C, “Familiar” sub-panels) and power spectra of LFP oscillations in the 1-40 Hz range (Fig. S8D,E, “Familiar” sub-panels). Upon switching to the novel environment, the *Cacna1c*^+/+^ rats had theta-gamma PAC levels similar to the initial response to the subsequently familiar environment (Fig. 4B,C, “Novel” sub-panels). However, the *Cacna1c^+/-^* animals showed a significantly impaired theta-gamma PAC compared to wild type animals (Fig. 4B,C).

**Fig. 4.**
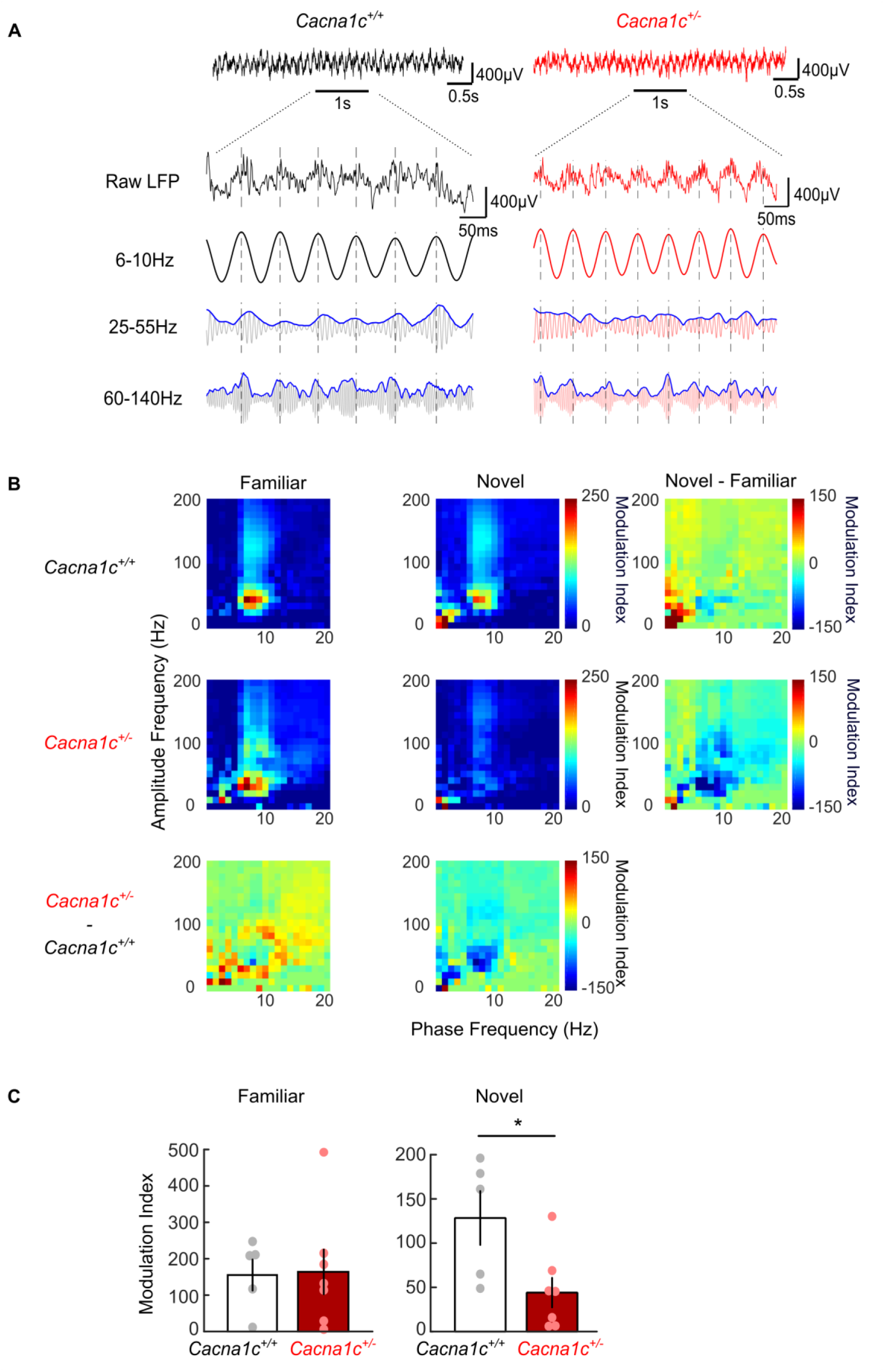
Reduced Theta-Slow Gamma Coupling in a Novel Environment in Cacna1c^+/-^ rats. (A) Local field potentials taken from runs on the familiar track. *Top*: Representative broadband LFP traces (0.1-475Hz) taken from dorsal CA1 in a 5 second window while running. *Bottom*: 1 second expanded segment showing raw unfiltered LFP (top) and bandpass filtered theta (6-10Hz), slow gamma (25-55Hz) and fast gamma (60-140Hz) (bottom). Phase-amplitude coupling measures the degree to which the amplitude envelope (blue traces shown for slow and fast gamma) is modulated by the phase signal (theta filtered LFP). Black: *Cacna1c^+/+^*; red: *Cacnac1^+/-^*; dashed vertical lines mark timings of theta cycle peaks. (**B**) Mean Phase-Amplitude Coupling comodulograms taken from track runs on the familiar and novel track. Color represents the modulation index (MI) depicting the degree of coupling between the phase of frequencies (1-20Hz) on the x-axis and the amplitude of frequencies (1-200Hz) on the y-axis. Differences between environments and genotypes are shown next to the mean plots as labelled. (*Cacna1c^+/+^*: n = 5; *Cacna1c^+/-^*: n=7). (**C**) Mean Phase-Amplitude Coupling between theta (6-10Hz) and slow gamma (25-55Hz). Theta-Slow Gamma coupling was lower in *Cacna1c^+/-^* rats than *Cacna1c^+/+^* rats on the novel track (p=0.0256; student’s t-test) but not on the familiar track (p=0.9184; student’s t-test). (*Cacna1c^+/+^*: n=5; *Cacna1c^+/-^*: n=7; Data presented as means ±SEM. *p <0.05.

Our results indicate that *Cacna1c* hemizygosity results in altered network properties in the hippocampus including a deficit of coupling between CA3 and CA1 subfields. This effect may disrupt hippocampal network physiology during the encoding of contextual information and contribute to the observed deficit in LI seen in *Cacna1c^+/-^* animals.

### Hippocampal levels of phosphorylated ERK and phosphorylated CREB are reduced in the Cacna1c^+/-^ rats

We next investigated molecular changes associated with altered synaptic plasticity and network activity in the hippocampus of *Cacna1c^+/-^* animals. Our observation of impaired spine Ca^2+^ signaling at SC-CA1 synapses in *Cacna1c^+/-^* rats suggests that low dosage of *Cacna1c* may impact calcium-dependent signaling pathways downstream of Ca_V_1.2 VGCCs. In particular, the ERK signaling pathway is known to couple L-VGCC activation to changes in CREB activation and gene expression required to support long-lasting plasticity (*30, 31*). We therefore examined (basal) levels of ERK and CREB and the levels of phosphorylated ERK (pERK) and phosphorylated CREB (pCREB) in the dorsal hippocampus using immunohistochemistry. We found a significant reduction in pERK (Figs. 5A, S9A) in all hippocampal subfields and reduced pCREB in CA1 and CA3 (Figs. 5B, S9C) in the *Cacna1c*^+/-^ compared to *Cacna1c*^+/+^ animals with no differences in overall ERK and CREB protein levels between the two genotypes (Fig. S9B,D). This marked decrease of hippocampal pERK and pCREB levels in *Cacna1c*^+/-^ rats indicates ERK pathway activation and subsequent nuclear signaling are highly sensitive to Ca_V_1.2 levels.

**Fig. 5.**
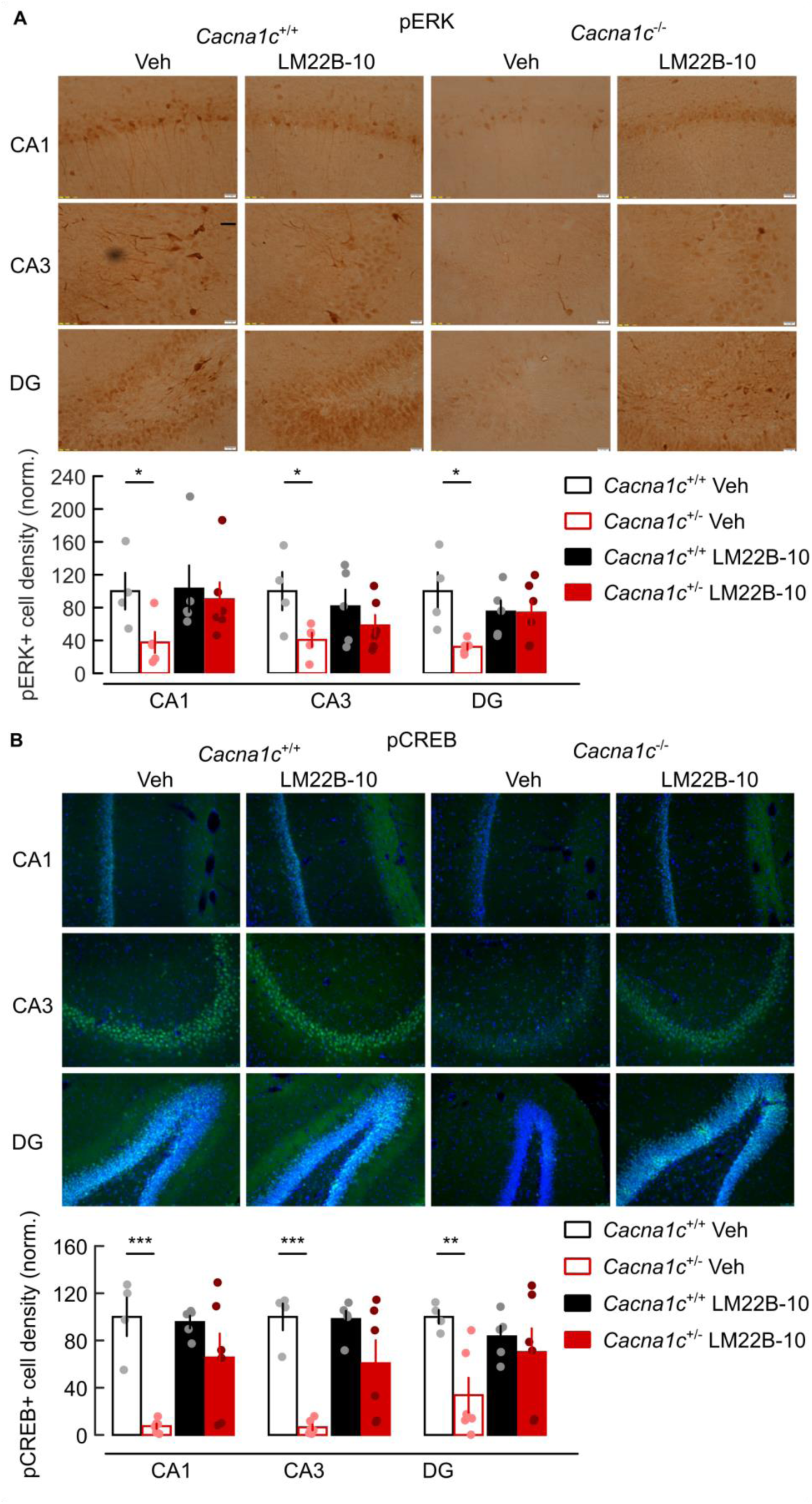
TrkB/TrkC receptor agonist LM22B-10 restored the levels of hippocampal pERK and pCREB in *Cacna1c^+/-^* rats. (**A**) Systemic administration of the TrkB/TrkC agonist LM22B-10 (25mg/kg, i.p.) restored the baseline hippocampal levels of pERK in *Cacna1c^+/-^* rats. Genotype × treatment interaction: F_(1,16)_=4.567, p=0.048; post-hoc comparison between *Cacna1c^+/+^ versus Cacna1c^+/-^* received vehicle (Veh) (F_(1,16)_=12.866, p=0.002) and those that received LM22B-10 (F_(1,16)_=0.615, p=0.444). (**B**) Systemically administered LM22B-10 restored the baseline level of hippocampal pCREB in *Cacna1c^+/-^* animals. Genotype × treatment: F_(1,17)_=5.239, p=0.035; post-hoc comparison between *Cacna1c^+/+^ versus Cacna1c^+/-^* rats that received vehicle (Veh) (F_(1,16)_=21.319, p=0.000) and those that received LM22B-10 (F_(1,16)_=2.485, p=0.133). All panels, *Top*: example immunoreactivity for pERK (A) and pCREB (B) in LM22B-10- or vehicle-treated animals; *Bottom*: summary of immunopositive cell densities in CA1, CA3 and dentate gyrus (DG) of dorsal hippocampus, normalized to average value in *Cacna1c^+/+^* tissue. Animals were sacrificed 60 min after i.p. administration of drug or vehicle. Sample sizes for *Cacna1c^+/+^*: LM22B-10 n=5, Veh n=4; *Cacna1c^+/-^*: LM22B-10 n=6, Veh n=5 (pERK) and n=6 (pCREB). Data presented as means ±SEM. *p <0.05, **p < 0.01 and ***p< 0.001, two-way repeated measures ANOVA followed paired wise post-hoc comparison with Bonferroni correction.

### Rescue of ERK signaling and associative plasticity in Cacna1c^+/-^ animals with TrkB/TrkC agonist LM22B-10

We hypothesized that pharmacological activation of the ERK pathway could reverse the molecular and physiological deficits observed in *Cacna1c^+/-^* animals. We targeted the ERK pathway by using a recently discovered small molecule BDNF mimetic TrkB/TrkC neurotrophin receptor co-activator (LM22B-10) previously shown to activate the ERK pathway signaling in neurons (*55*). Systemic administration of LM22B-10 (25 mg/kg i.p.) restored the baseline levels of pERK and pCREB activation in the dorsal hippocampus in *Cacna1c*^+/-^ rats to the levels seen in *Cacna1c*^+/+^ animals 60 min after injection (Fig. 5). LM22B-10 did not however alter the number of pERK- and pCREB-positive cells in wild-type hippocampus.

We next determined whether LM22B-10 could also rescue the hippocampal synaptic plasticity in *Cacna1c*^+/-^ rats. TBP-induced LTP at SC-CA1 synapses was rescued in *ex vivo Cacna1c*^+/-^ slices by LM22B-10, either bath-applied *in vitro* (Fig. 6A,D) or in *Cacna1c*^+/-^ slices prepared 60 min after systemic administration (Fig. 6B,D). LM22B-10 did not affect TBP-induced LTP in *Cacna1^+/+^* slices (Fig. 6C,D). These results show that the activation TrkB/TrkC receptors can bypass the Ca_V_1.2 L-type VGCC signaling cascade to rescue ERK signaling and associative hippocampal plasticity in *Cacna1c*^+/-^ rats.

**Fig. 6.**
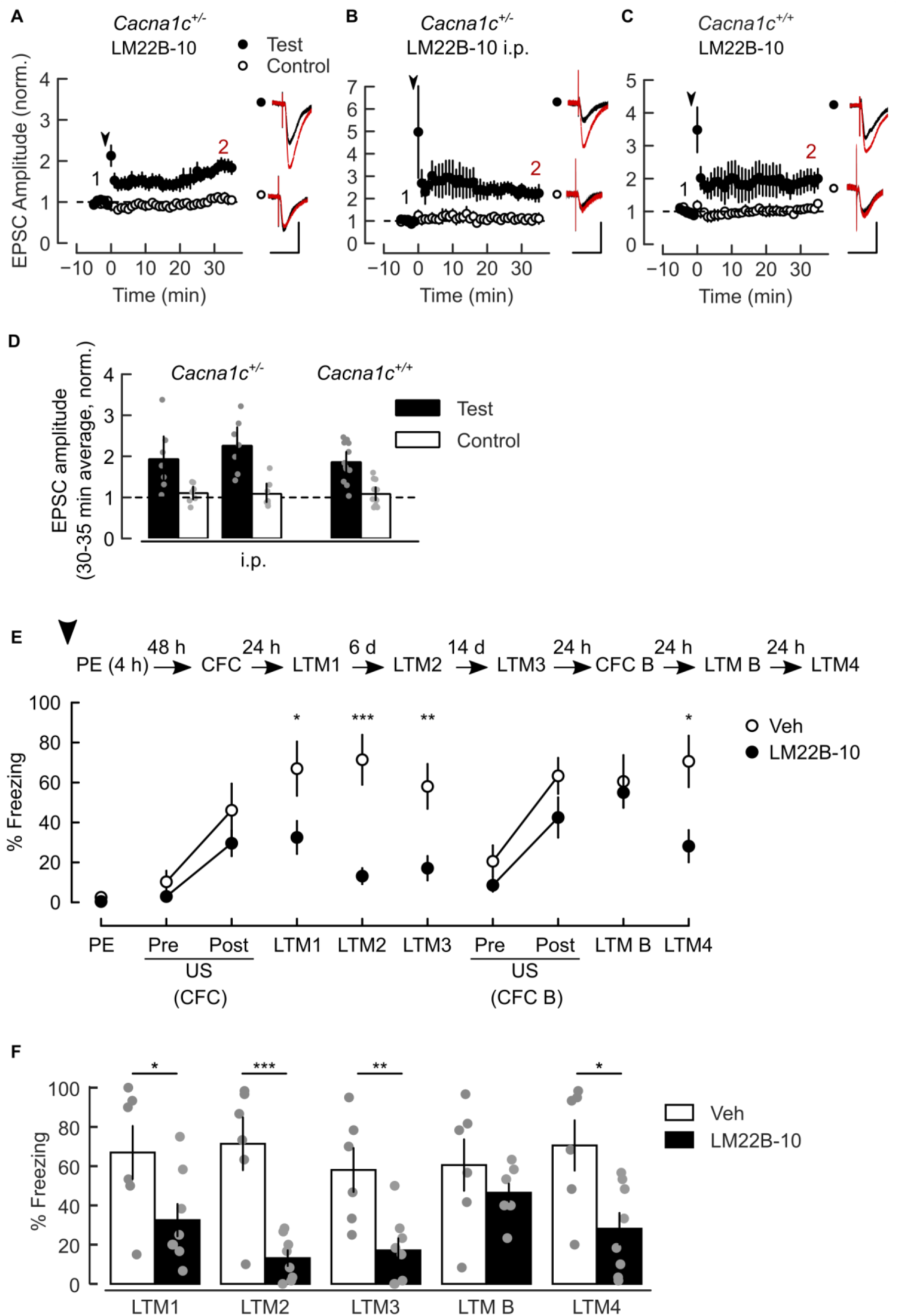
The TrkB/TrkC receptor agonist LM22B-10 rescued associative LTP at SC–CA1 synapses and LI of CFC in *Cacna1c^+/-^* rats. (**A,B**) The TrkB/TrkC agonist LM22B-10 rescued SC-CA1 LTP induction with TBP in slices from *Cacna1c^+/-^* rats, either during bath application (2 µM, A) or after i.p. bolus (25 mg/kg, B). (**C**) Bath-application of LM22B-10 did not affect TBP-induced SC-CA1 LTP in slices from wild-type animals. (**D**) Summary of change in normalized mean EPSC amplitude at 30-35 min after LTP. Pairwise Test vs Control, A: p=0.034, n=12/4; B: p=0.032, n=7/4; C: p=0.044, n=7/4. Sample sizes given as cells/animals. Schematic of TBP protocol given in Fig. 3A. (**E**) Intrahippocampal infusion of LM22B-10 before context pre-exposure rescued LI of CFM in *Cacna1c^+/-^* animals. *Schematic*: the behavioral paradigm of Fig. S1 extended with CFM retrieval trials in LI context (LTM3, LTM4), training in alternative context (CFC B) and retrieval in alternative context (LTM B). Arrowhead: bilateral infusion of 1µl LM22B-10 (2 µM, n=8) or vehicle (Veh, n=6) into the dorsal hippocampus 60 min before PE. The plot shows the freezing response in all sessions. (**F**) Summary of freezing response at LTM tests. The freezing responses of LM22B-10- and vehicle-infused animals were different (treatment × stage interaction: F_(5,60)_=5.205, p=0.002). Compared to the vehicle group, LM22B- 10 infused animals showed reduced freezing responses only in the LI context (between-subject contrasts *versus* Veh, LTM1: F_(1,11)_=4.162, p=0.033, LTM2: F_(1,11)_=18.467, p=0.0005, LTM3: F_(1,11)_=9.602, p=0.005, LTM4: F_(1,11)_=6.886, p=0.012, LTM B: F_(1,11)_=0.138, p=.358). Data presented as means ±SEM. *p <0.05, **p < 0.01 and ***p<0.001, two-way repeated measures ANOVA followed by pairwise post-hoc comparison with Bonferroni correction.

### TrkB/TrkC agonist LM22B-10 rescues the behavioral deficits in Cacna1c^+/-^ rats

Given that LM22B-10 reverses the molecular and plasticity deficits in *Cacna1c*^+/-^ rats, we investigated whether TrkB/TrkC activation in the hippocampus could rescue the behavioral deficit in LI of contextual fear memory observed in these animals. To explore this possibility, we initially directly infused LM22B-10 into the dorsal hippocampus during the pre-exposure stage of LI training in *Cacna1c*^+/-^ rats. A single infusion of LM22B-10 into the dorsal hippocampus 60 min prior to the pre-exposure rescued the LI effect when animals were tested 24 h, 7 days and 21 days after CFC training (Fig. 6E). To assess whether LM22B-10 infusion had non-specific confounding effects on hippocampal function, all rats received a second CFC in a separate novel context (context B) in the absence of any infusion 22 or 23 days later. All rats expressed equivalent levels of freezing responses to context B and normal hippocampal-dependent CFC and fear memory in the novel context (Fig. 6E,F). In addition, when LM22B-10 infused rats returned to the original conditioned context at LTM4 they showed intact and context specific LI of CFC. Thus, intrahippocampal infusions of LM22B-10 rescued LI without producing any deleterious effects on hippocampal function.

We further investigated whether intra-peritoneal administration of LM22B-10 could rescue the behavioral deficits in *Cacna1c*^+/-^ rats. Our results show that systemic injections of LM22B-10 through the LI procedure can also rescue the LI effect in *Cacna1c*^+/-^ rats (Fig. S10). Together, these findings show that activation of the ERK signaling pathway in the dorsal hippocampus during the pre-exposure phase of LI is sufficient to rescue the behavioral deficits seen in *Cacna1c*^+/-^ rats. Furthermore, this effect can be achieved with both intrahippocampal administration of LM22B-10 and with systemic dosing.

## Discussion

The past decade has seen major advances in psychiatric genomics. These provides grounds for optimism that greater mechanistic insight can be gained into disorders such as schizophrenia and bipolar disorder facilitating the development of new therapies. The consistent association of schizophrenia, bipolar disorder and related neurodevelopmental disorders with genetic variation in VGCCs, and in particular *CACNA1C*, makes understanding the impact of genetic variation in the associated loci of high importance in terms of advancing mechanistic understanding of these conditions. Here we show that reduced dosage of *Cacna1c*, which is associated with neurodevelopmental and psychiatric disorders in humans, produces a distinctive behavioral, physiological, and molecular phenotype in rodents. We also show that the impact of low dosage of *Cacna1c* on behavior, plasticity and signaling pathways can be reversed by the intracerebral or peripheral administration of a small molecule drug targeting the Trk family of neurotrophin receptors, highlighting a novel target for therapy in these conditions.

Cognitive abnormalities have been consistently associated with a range of psychiatric presentations. Psychotic symptoms, which are pathognomonic of schizophrenia and are frequently seen in bipolar disorder, have been associated with altered learning about the associations between environmental stimuli (*16, 17, 19, 21*). Such altered associative learning can result in aberrant attribution of importance (or salience) to otherwise irrelevant events (*56*). One measure of such altered learning, which has been shown to be affected in individuals with psychosis, is the latent inhibition procedure (*15*). Here we found that low dosage of *Cacna1c* was sufficient to produce a marked deficit in contextual latent inhibition consistent with the impairments seen in psychotic patients. This deficit was seen despite *Cacna1c* hemizygous animals showing normal contextual fear conditioning. These results are consistent with previous studies of mice with homozygous genetic deletions of *Cacna1c* (which encodes the pore-forming subunit of Ca_V_1.2 L-VGCCs) or *Cacna1d* (which encodes the pore-forming subunit of Ca_V_1.3 L- VGCCs) which have shown that Ca_V_1.3, but not Ca_V_1.2, is essential for the acquisition of fear associations (*35, 57*). In contrast there is evidence from brain-specific knock-out studies that Ca_V_1.2, but not Ca_V_1.3, is required for context discrimination (*25, 58*). This suggests that the encoding of the contextual information required to mediate latent inhibition may be particularly susceptible to alterations in the dosage of *Cacna1c*.

The generation of novel context representations is dependent on the hippocampus and involves the recruitment of CA1 pyramidal neuronal ensembles, or engrams, driven by the conjunctive excitatory afferents from the entorhinal cortex and CA3 hippocampal area (*50, 52*), and supported by synaptic plasticity induced during theta burst firing events (*59, 60*). We hypothesized that in *Cacna1c^+/-^* rats, the low Ca_V_1.2 dosage in CA1 pyramidal neurons in the dorsal hippocampus would impact on both synaptic plasticity at CA1 synapses and on the rhythmic coordination of synaptic activity in the CA1 area.

Associative synaptic potentiation can be induced by temporally coordinated pre- and postsynaptic neuronal activity and requires the activation of postsynaptic NMDARs and VGCCs (*26, 27, 42*). We found that induction of associative long-term potentiation (LTP) by coincident pre- and postsynaptic spikes (TBP-LTP) is impaired at SC-CA1 synapses in *Cacna1c*^+/-^ animals. However, the SC-CA1 synapses in *Cacna1c*^+/-^ rats appear mechanistically capable of plasticity and can be strengthened under conditions not requiring the activation of L-VGCCs as shown by using an LFS-pairing protocol known to induce NMDAR-dependent but L-VGCC-insensitive LTP (*43*). In contrast to the loss of TBP-LTP at SC-CA1 synapses, induction of associative plasticity was notably preserved at TA-CA1 synapses, which represent the other major source of cortical excitatory inputs onto CA1 pyramidal neurons in the hippocampus.

In order to further investigate the causes of this selective loss of TBP-LTP at SC-CA1 synapses in the hippocampus we investigated the excitability of, and calcium signaling in, CA1 pyramidal neurons in *Cacna1c*^+/-^ rats. *Cacna1c*^+/-^ neurons showed a reduction in the frequency-dependent broadening of somatic action potentials (APs) during repetitive discharge. This effect was significant at firing frequencies that occur during LTP induction with TBP and was replicated in wild-type cells by application of isradipine, confirming a role for L-VGCCs. Narrower somatic APs elicit weaker dendritic Ca^2+^ signals in CA1 pyramidal neurons, when compared to broader APs (*47*). Our Ca^2+^ imaging experiments reveal an inefficient summation of spine Ca^2+^ signals during bursts of back-propagated APs at SC-CA1 synapses in *Cacna1c*^+/-^ neurons. The reduced somatic AP broadening and the diminished spine Ca^2+^ signals during AP bursts indicate that the low *Cacna1c* dosage impacts on the association between postsynaptic spiking and synaptic input necessary for associative plasticity to occur at SC-CA1 synapses (*27, 44*). The deficit in action potential-triggered Ca^2+^ signaling was significant in dendritic spines in the *stratum radiatum*, where the majority of Schaffer collateral (SC) synapses are made (*61*). Our findings that TA-LTP is intact in *Cacna1c^+/-^* neurons suggest that the reduced *Cacna1c* dosage does not affect L-VGCC-dependent mechanisms in the apical tuft, where local dendritic spikes appear more important for LTP induction (*26*). We propose that the selective impairment of forms of LTP that require post-synaptic spiking in addition to NMDAR activation at SC-CA1 synapses underpins the selective behavioral phenotype seen in *Cacna1c*^+/-^ animals.

Network synchronization, and in particular theta-gamma PAC, in area CA1 are thought to contribute to the establishment of hippocampal memory traces by providing adequate temporal coordination between pre-and postsynaptic spiking (*22*). We found a deficit in the phase-amplitude coupling between theta and slow-gamma oscillations in the CA1 area of *Cacna1c^+/-^* rats, manifested specifically during the exploration of a novel environment, indicating an impaired coupling between CA1 neurons and excitatory input from area CA3 (*52*). Alterations in theta-gamma PAC across brain regions has been recently reported in animal models of psychosis (*23, 62*) and in schizophrenic patients (*63, 64*), and therefore may reflect a broader spectrum of network deficits in psychoses. Taken together these results show that low dosage of *Cacna1c* impairs both plasticity and network activation in the hippocampus in a manner likely to contribute to the observed deficits in contextual learning and latent inhibition seen in *Cacna1c^+/-^* rats.

At a molecular level L-VGCC-mediated Ca^2+^ signaling is essential for excitation-translation coupling. Ca^2+^ influx via L-VGCC imparts location and temporal specificity for activation of the ERK signaling pathway(*30*), necessary for mechanisms of early LTP including AMPA receptor trafficking (*28, 29*) and for protein synthesis-dependent forms of long-term potentiation and memory (*65*). Here we show that *Cacna1c*^+/-^ rats have markedly decreased levels of ERK and CREB activation as assessed by their phosphorylated levels in the hippocampus. To test the hypothesis that impaired ERK signaling downstream Ca_V_1.2 contributes to the behavioral and hippocampal functional deficits in *Cacna1c*^+/-^ animals, we sought to activate the ERK pathway independently of L-VGCCs. Recent evidence supports the targeting of neurotrophin receptors using small molecule BDNF mimetics as potential therapeutic strategy in a range of neuropsychiatric and neurodegenerative disorders (*66, 67*). With a view to rescuing synaptic dysfunction and spine deficits in the Cacan1c+/- animals, we used a small molecule TrkB/TrkC co-activator LM22B-10, previously shown to activate the ERK pathway in mice *in vivo* (*55*).

Direct application of LM22B-10 on slices rescued TBP-induced LTP at SC-CA1 synapses in *Cacna1c*^+/-^ rats, and direct hippocampal infusion of LM22B-10 during context preexposure was sufficient to rescue the observed behavioral deficit in LI. In addition, we were able to show that peripheral (systemic) treatment with LM22B-10 was sufficient to rescue the observed molecular and plasticity changes in the hippocampus and to restore normal LI behavior in *Cacna1c*^+/-^ rats. These results suggest that the targeting of neurotrophin receptor pathways may represent a route to ameliorating the deficits produced by altered dosage of *CACNA1C* in neuropsychiatric disorders.

In conclusion by adopting a genomically informed approach to investigate pathological processes associated with genetic risk for neuropsychiatric disorders we highlight the potential of drugs targeting neurotrophin receptor signaling as novel therapeutics in major psychiatric disorders including schizophrenia and bipolar disorder.

## Materials and Methods

### Study design

We used a *Cacna1c^+/-^* rat model in a multi-disciplinary, controlled laboratory experiment design with the objective to characterize the impact of reduced dosage of the psychiatric risk gene *CACNA1C* on behavior, synaptic and circuit function and to explore ways to rescue the observed deficits. LI of CFC was chosen as the central behavioral paradigm of the study because it explores hippocampal-dependent associative learning implicated in cognitive functions translationally relevant to psychoses (*15-17, 21, 68*). The *in vivo* and *ex vivo* electrophysiology and two-photon imaging experiments in the dorsal hippocampal CA1 subfield were informed by the observed context-sensitive behavioral deficits in *Cacna1c^+/-^* rats. These experiments were designed to investigate the impact of *Cacna1c* heterozygosity on hippocampal neural circuit synchronization, synaptic plasticity, excitability, and spine Ca^2+^ signaling as potential mechanisms underlying the behavioral deficit. The immunohistochemistry experiments were conducted to determine the impact of reduced *Cacna1c* dosage on the ERK/CREB signaling pathways downstream L-type VGCCs which has been implicated in synaptic plasticity, associative learning, and context-dependent learning(*28, 30, 65*). For the rescue experiments we administered a small molecule TrkB/TrkC receptor agonist previously characterized for ERK pathway activation activity and neurotrophic effects after systemic administration (*55*) either intrahippocampally or systemically (intra-peritoneal injections). When given peripherally, the agonist (or vehicle) was administered prior to each behavioral manipulation to account for non-specific state dependent effects (*69*).

### Animals

*Cacna1c* hemizygous rats with Sprague-Dawley background (*33*) were generated from cryo-preserved embryos (strain SD-Cacna1ctm1Sage-generated using zinc-finger nuclease technology, Sage Research Labs, Pennsylvania, USA) for a truncating mutation in exon 6 of *Cacna1c* gene. Animals were bred at Charles River (Margate, UK), Cardiff and Bristol Universities. We used a total of 278 *Cacna1c* wild type and heterozygous littermates. Behavioral experiments in Fig. S1 used 54 male Lister Hooded rats (250-275g, Charles River, UK). Littermates were housed in groups of up to four animals per group, with access to food and water *ad libitum*. Animals used in behavioral and immunohistochemistry studies were housed under reversed light/dark cycle (lights out between 10 a.m. - 8 p.m.) and behavioral training began at approximately 11 a.m. on each day. All other animals were maintained under normal 12h/12h light cycle. *Ex vivo* electrophysiology and two-photon imaging experiments were performed on 3-6 months old animals. *In vivo* electrophysiology recordings were made in 21-53 weeks old animals. All animal handling was performed in accordance with Home Office regulations and as directed by the Home Office Licensing teams at Cardiff and Bristol Universities.

### Statistical analysis

Sample sizes were determined by power analysis based on effect sizes routinely observed in the laboratory. Animals were assigned pseudo-randomly to each experimental group. Experimenters were blind to the genotype of animals when collecting the data. Behavioral and immunohistochemistry data was analyzed with repeated measures analysis of variance (ANOVA). Sphericity was tested using Mauchly’s Test and the Greenhouse–Geisser correction was applied if necessary and reported. Sources of significance were determined post-hoc using Tukey’s test. When drugs were used, only post treatment data was included in the ANOVA. Pairwise comparisons with Bonferroni correction were performed when ANOVA showed significant interaction between factors. In Fig. S9 two statistical outliers (> ± 2 SD) were removed from the analysis of immunohistochemistry data for basal phosphorylated CREB and phosphorylated ERK (one *Cacna1c*^+/-^ rat each). For synaptic plasticity experiments, comparisons were made between normalized mean EPSC amplitudes at 30-35 min after conditioning, with Test/Control pathway (within subjects factor) and genotype or drug treatment (between subjects factor). The effect of genotype or drugs on LTP induction were tested for significance using a two-way ordinal regression (cumulative link model) followed by analysis of deviance (ANODE). Passive membrane properties were compared using two-sample Mann-Whitney U test between genotypes against the null hypothesis of no difference between genotype-specific means. AP parameters for the first five or 10 APs during spike train discharges were compared using two-way ordinal regression with repeated measures for AP number and factors of genotype or first ISI frequency calculated as the reciprocal of first inter-spike interval. AP-elicited spine Ca^2+^ transients were compared within somatic distance zones using a generalized linear model with factors of genotype and number of APs, followed by analysis of variance (ANOVA). Tukey-adjusted p-values for post-hoc pairwise comparisons are indicated in figure legends together with the likelihood ratio chi square test values (LR), F statistic, and degrees of freedom for main effects and interactions. For *in vivo* electrophysiology, group comparisons were carried out using student’s t-test. Pooled data in all figures are represented as mean ± standard error of the mean (SEM). Values are presented as mean ± SEM. Statistical significance was established at p < 0.05. Throughout the analysis, the level of significance was set to 0.05 and the calculated probabilities are symbolized by asterisks as follows: * p < 0.05; ** p < 0.01; *** p < 0.001.

Detailed descriptions of the experimental procedures are provided in Supplementary materials.

## Acknowledgments

We thank J Carter, I Morella and C Best for technical support. We thank Prof. JR Mellor and Prof. R Brambilla for helpful discussions.

## Author contributions

C.M.T, T.E.L., L.S., M.C.O, M.J.O., L.S.W, E.M., M.W.J., K.L.T. and J.H. designed the experiments and analyses, and wrote the manuscript. T.E.L., L.S. and K.L.T. performed behavioral experiments. C.M.T. performed *ex vivo* electrophysiology and imaging experiments. E.M. performed *in vivo* electrophysiology experiments. T.E.L. and K.L.T. performed the molecular studies.

## Supplementary Materials

### Materials and methods

#### Behavioral paradigms

##### Contextual fear conditioning (CFC), contextual fear memory (CFM)

A paradigm of a single footshock exposure in a novel context was used (*70*). During 3-min CFC training trial, each animal received a single scramble footshock (US: 0.5mA, 2s) after being placed in a conditioned chamber (context) for 2 min. Animals were returned to home cage after conditioning. CFM retrieval was tested by measuring the animal’s freezing response upon returning to the conditioned chamber for 2 min. Retrieval tests were performed 3 h (short-term memory, STM), 24 h (long-term memory LTM1), 7 days (LTM2) and 21 days (LMT3) and 23 days (LTM4) after CFC training. Intrahippocampal LM22B-10 infusion rats received additional CFC training and CFM retrieval in an alternative context (context B) on Days 22 and 23 after the first CFC training. The freezing behavior was used as measure of conditioned fear. One unit of freezing response was defined as the continuous absence of movement other than respiratory motion in 1 s sampled every 10 s. Freezing responses were video-recorded during each trial and quantified as percentage of time spent freezing by an observer blind to the experimental group.

##### Latent inhibition of contextual fear conditioning (LI of CFC)

LI training consisted of animal preexposure (4 h, PE) to the conditioned context, either 24h or 48 h prior to CFC training.

##### Apparatus for behavioral testing

All sessions were undertaken in standard modular test chambers for rats (interior dimensions: 30.5 cm x 24.1 cm x 21.0 cm) placed inside sound attenuating chambers 55.9 cm x 55.9 cm x 35.6 cm (Med Associates Inc., Vermont, USA). Two test chambers were used as a standard context for single CFC, one consists of standard transparent Perspex walls, and the other chamber was decorated with black and white stars wallpaper on the Perspex. Both chambers were equipped with 19 equally spaced metal bars placed 1.6 cm above the floor, for the delivery of a 0.5 mA current (footshock) under control by a stand-alone aversive stimulator/scrambler (Med Associates Inc., Vermont, USA). Rats were placed in either of the chamber for single CFC training. The chambers were wiped with 50% EtOH prior placing the animals in the chamber. For rats that received additional CFC training in an alternative context, had they received original CFC in transparent Perspex chamber (context A) would be placed in star decorated chamber, wiped with 5% pepper mint water (context B) for the second CFC training. The programmes for each session were controlled through a Med-PC version IV research control and data acquisition system (Med Associates Inc., Vermont, USA). Behavior was video recorded with cameras (JSP Electronics Ltd, China) positioned centrally above the chambers, digitized and analyzed offline using Numeroscope software (Viewpoint, France). For each training or retrieval session, animals were individually transferred between from home cages and the testing room in the same large transport box.

#### In vivo electrophysiology

##### Surgical Procedure

All surgery was carried out aseptically to minimize the risk of infection. Animals were anaesthetized with isoflurane in oxygen (maintained at 1.5-2%) and head-fixed in a stereotaxic frame (Kopf model 1900). After exposing the skull surface, a 2.5mm craniotomy was made using a tungsten carbide burr (3.6mm posterior to bregma and 2.5mm lateral to the midline) and a 20-tetrode (16 recording electrodes and 4 reference electrodes) microdrive was lowered into the craniotomy (Fig. S8A). The microdrive was fixed in place using dental cement (dePuY). Animals were given buprenorphine (0.025mg/kg) following surgery, and weight and water consumption were monitored for at least one week afterwards. Over a duration of 2-4 weeks tetrodes were gradually lowered (∼20-40µM a day) to the pyramidal cell layer of CA1 (verified by the presence of sharp-wave ripples and bursting spike activity).

##### Data Acquisition

Recordings were made using a Digital Lynx SX system (Neuralynx). Local field potentials were sampled at 1kHz and filtered between 0.1 and 475Hz). Animal position was monitored with an overhead camera and an LED attached to the headstage.

##### Training

Animals were trained to run back and forth on a 175cm linear track over a period of 1-4 weeks. During this period animals were food restricted to 85% of their initial weight. Behavioral experiments were carried out once tetrodes were in place in the pyramidal cell layer of dorsal CA1 and animals had reached a minimum criterion of 10 complete circuits on the linear track.

##### Familiar Track Behavioral Protocol (Fig. S8B)

Animals were placed in a sleep-box (a sound-attenuating chamber) and left to rest for approximately 30 minutes prior to track running. After the pre-sleep period animals were allowed to run freely for sucrose rewards. Following the run, animals were placed back in the sleep-box for a further 30 minutes. Animal position and electrophysiological signals were continuously recorded throughout the behavioral session.

##### Novel Track Behavioral Protocol (Fig. S8C)

Following at least one recording on the familiar track and at least a day after the familiar track session, animals were subject to the novel position behavioral protocol. The pre and post sleep protocol here did not differ to that of the familiar track behavioral sessions. For the track sessions animals were placed on the familiar track for approximately 10 minutes to take baseline recordings. The animals were then taken off the track, the track was rotated 45° and animals were immediately placed back on and recorded while they now freely ran through a different portion of the recording room.

##### Data Analysis

All data analysis of in vivo electrophysiology was carried out in MatLab (Mathworks). Local field potential signals were taken from periods of track running over 35cm/s and all analysis was performed on these segments of local field potential. To control for individual animal differences in the number of circuits on the familiar track compared to the novel track, a matched number of runs were pseudo-randomly selected from the session containing more runs.

##### Power Spectral Analysis

All power spectral estimates were calculated using multi-taper spectral analysis (Chronux toolbox) with a window length of 2.5s, a time-bandwidth product of 3 and 5 tapers.

##### Phase-Amplitude Coupling

Phase-amplitude coupling was detected using a PAC toolbox applying the modulation index measure. (Onslow et al., 2011). In brief this method combines the amplitude envelope of the local field potential filtered at a high frequency band with the phase values of the signal filtered at a lower band to form a composite signal. The mean of this signal provides a measure of the strength of coupling between the amplitude of the higher frequency band with the phase of the lower frequency band.

#### Ex vivo electrophysiology

##### Slice preparation and recordings

Acute transverse hippocampal slices were obtained from 3 – 6 months old male and female SD-Cacna1c rats after being given a lethal dose of isoflurane inhalation. The hippocampi were dissected in ice-cold slicing solution (in mM: 110 Choline chloride, 25 glucose, 25 NaHCO_3_, 2.5 KCl, 1.25 NaH_2_PO_4_, 0.5 CaCl_2_, 7 MgCl_2_, 11.6 L-Ascorbic acid, 3 pyruvic acid), then mounted on agar blocks and cut in 400 µm thick slices on a Microm HM 650V vibratome (Thermo-Scientific). Slices were incubated in recovery solution (in mM: 93 N-methyl-D-glucamine, 25 glucose, 30 NaHCO_3_, 20 HEPES, 2.5 KCl, 1.2 NaH_2_PO_4_, 0.5 CaCl_2_, 7 MgCl_2_, 5 L-Ascorbic acid, 3 pyruvic acid, 12 N-acetyl-cysteine, 2 thiourea) at 35 °C for 5 min, then incubated in artificial CSF (aCSF, in mM: 119 NaCl, 10 glucose, 26.2 NaHCO_3_, 2.5 KCl, 1 NaH_2_PO_4_, 2.5 CaCl_2_ and 1.3 MCl_2_), at 35 °C for 30 min. Slices were then stored in aCSF at room temperature until use. Solutions were equilibrated with 95% O_2_ and 5% CO_2_ and had osmolarity of 300 – 310 mOsm. Whole-cell patch-clamp recordings were made from CA1 pyramidal neurons visualized under differential interference contrast (Olympus BX-51 WI) in a submerged recording chamber superfused with aCSF (∼ 2 ml/min) at 35 °C. Patch electrodes (4 – 6 MΩ) were pulled from borosilicate filamented capillaries on a Model P-1000 Flaming/Brown micropipette pulse (Sutter Instruments) and filled with intracellular solution (in mM: 117 KMeSO_3_, 8 NaCl, 1 MgCl_2,_ 10 HEPES, 4 MgATP, 0.3 NaGTP, pH 7.2, 280 mOsm, 0.2 EGTA). For the experiments in Fig 2f-k involving tonic postsynaptic depolarization the firing of action potentials was blocked by substituting CsMeSO_3_ for KMeSO_3_ and adding 5 mM QX-314 to the internal solution (pH was adjusted with CsOH). Recordings were made with a Multiclamp 700B amplifier and digitized with an Axon Digidata 1550 Data Acquisition System and pClamp 10.7 software (Molecular Devices).

##### Synaptic plasticity experiments

Synaptic currents were recorded in voltage-clamp (−70 mV), and the aCSF was supplemented with 50 µM picrotoxin to block GABA_A_ receptors. Synaptic responses were evoked with 0.1-1 ms square pulses delivered through bipolar tungsten electrodes (100 kΩ, MicroProbes) placed on opposite sides of the recorded cell, in *stratum radiatum* (schematic in Fig 2A,E) or in *stratum lacunosum-moleculare* (schematic in Fig 2I). For synaptic plasticity experiments synaptic currents elicited at 0.1 Hz alternatively in Test and Control pathways were filtered at 4 kHz and digitized at 10 kHz. When used, back-propagated postsynaptic action potentials (bAPs) were evoked by somatic current injection (2 nA, 2 ms).

Membrane voltage was not corrected for liquid junction potential, calculated at −9 mV. The pathways were tuned in current-clamp to evoke sub-threshold EPSPs prior to baseline recording and were tested for independence using paired-pulse protocols. Data was analyzed offline using routines written in Python. Consecutive excitatory post-synaptic currents (EPSC) in each pathway were averaged every minute and their amplitudes normalised to the average of 5 min before the application of a conditioning protocol to the Test pathway (baseline), while the Control pathway was left unperturbed. The mean EPSC amplitude in the Test and Control pathway at 30 – 35 min after conditioning was normalized to the mean EPSC amplitude during the 5 min baseline. The theta-burst pairing (TBP, Fig. 2A-D and I-L, S3) conditioning protocol consisted of three 1s trains of short pre- and postsynaptic bursts delivered at 5 Hz every 10s intervals. Each burst consisted of five of coincident pre-synaptic stimuli and post-synaptic bAPs at 100 Hz. The low-frequency stimulation pairing (LFS-Pairing, Fig. 2E-Hprotocol consisted of a train of pre-synaptic stimulations delivered at 2 Hz for 90 s, paired with tonic postsynaptic depolarization at 0 mV. Series resistance was monitored throughout the recording and cells with series resistance > 30 MΩ or showing > 20% change in series resistance were discarded. When used, drugs were continuously in the perfusate throughout the experiment.

##### Passive and active electrophysiological membrane properties

Membrane properties were determined from membrane potential (V_m_) recordings during 500 ms square-wave current injections in whole-cell current clamp, using the amplifier bridge circuit (*71*). The recordings were filtered at 6 kHz and digitized at 50 kHz. After determining the resting membrane potential (RMP), the membrane potential was set to −70 mV by a small inward current injection. The membrane time constant (τ_m_), the hyperpolarization-activated “sag” potential (V_sag_) and the input resistance (R_in_) were determined current injections of −0.1 nA (schematic in Fig S4A). V_sag_ was measured as the difference between the V_m_ minimum and the steady-state hyperpolarization elicited by the injected current, and expressed as a percentage of the steady-state membrane voltage (V_ss_); τ_m_ was determined from an exponential fit of the V_m_ curve between 10% – 95% from baseline to the V_sag_ minimum; R_in_ was calculated according to the Ohm’s law (V=I×R) using the difference between the steady-state hyperpolarization (V_ss_) and the resting membrane potential (RMP). Active membrane properties were determined from V_m_ recordings during a series of depolarizing current injections steps from 0 to +0.7 nA in 50 or 100 pA increments every 10 s, to elicit action potential (AP) firing (Fig. S4B). The rheobase current (I_rh_) was determined using exponential fits through strength-latency curves representing the injected current intensity as a function of spike latency (*72*). AP waveforms were detected by thresholding the first-order derivative of the recordings, and V_m_ onset was determined as the membrane voltage where the rate of change of membrane potential (dV/dt) exceeded 20 V/s (*73*). AP waveform duration was measured at one third from onset to peak, and the maximum rate of change (dV/dt) of the V_m_ during an AP was determined from the first-order derivative of the AP waveform. The instantaneous AP was calculated as the inverse of the inter-spike interval (ISI). To compare the relationship between AP durations and the firing rate, the AP trains recorded in each cell were grouped according to the instantaneous frequency at the 1^st^ ISI, in four frequency bands (Fig. S4F-I) and the durations of individual APs were averaged across the trains in the same frequency band. Some cells did not fire AP trains with frequencies falling in all four bands. The number of cells which fired AP trains in the given frequency band are indicated in the legend of Fig. S4. When used, isradipine was added to the perfusate following an initial series of current injections (“Vehicle”), and recordings were restarted after 7-10 min of drug wash-in.

#### Two-photon spine Ca^2+^ imaging

##### Data acquisition

Spine Ca^2+^ imaging was performed on a galvanometer-based two-photon laser scanning system (Ultima In Vitro Multiphoton Microscope System with Prairie View v.5.0 software, Bruker France – Nano Surfaces) in CA1 pyramidal neurons in *ex vivo* transverse hippocampal slices. Patch electrodes were filled with intracellular solution (in mM: 117 KMeSO_3_, 8 NaCl, 1 MgCl_2,_ 10 HEPES, 4 MgATP, 0.3 NaGTP, pH 7.2, 280 mOsm) supplemented with the medium affinity fluorescent Ca^2+^ indicator (Fluo-5F 200 µM; Life Technologies) and a reference fluorescent dye (AlexaFluor 594, 30 µM; Life Technologies). EGTA was omitted to avoid introducing additional Ca^2+^ buffering capacity in the cells. Fluorescence was excited with a Chameleon Ultra II Ti:Sapphire laser (Coherent) tuned at 810 nm. Ca^2+^ transients elicited by back-propagated action potentials (APCaTs) were imaged in dendritic spines in the *stratum radiatum* using line scanning mode (*74*) with a 60× water-immersion objective. After establishing whole-cell configuration in voltage-clamp, the cells were switched to current-clamp, the RMP was measured, and the membrane voltage was set to −70 mV by inward current injection (−50 pA – 100 pA). Cells were dye-loaded for 10-15 min then APCaTs were recorded in line-scanning mode, in series of 1000 lines per second, for 1s. To minimize photodamage, line scans were acquired in batches of up to 20 repeats every 20 s. APCaTs were evoked by delivering either one, two, three or five APs by brief (2 nA, 2 ms) somatic current injections (Fig. 3). Between batches, the RMP was monitored and cells where the RMP exceeded −60 mV or varied by more than 10 mV were discarded. Cells were discarded if they showed signs of photodamage (localized swellings, sustained increase in resting Ca^2+^ fluorescence, or tonic increase in Ca^2+^ fluorescence after stimulus, without return to baseline levels). Within the batches, the line scan series were interleaved with respect to the number of delivered APs, to compensate for small variations in dye loading between the beginning and the end of the imaging session. The trajectory of the line scan was set to image 3 – 10 spines per dendritic segment, on branches up to the third order in *stratum radiatum*. To minimize the risk of photodamage and surplus Ca^2+^ buffer capacity due to continuous dye accumulation we recorded spine APCaTs from up to three different dendritic locations and a single dendritic segment was imaged in each location. The total imaging time was limited to 50 – 60 min since break in. The locations of spines were selected pseudo-randomly with the constraint that the parent dendritic segments belong to distinct branches originating from the main trunk at least 30 µm apart (Fig. S5).

##### Data analysis

Fluorescence images were analyzed off-line using software written in Python. Line scan images were de-noised and data from up to five replicates in each protocol were averaged for each spine, including failures. The small number of replicates in each protocol precluded reliable calculation of success rates. APCaT traces were then generated as the relative change in Fluo-5F fluorescence channel *versus* Alexa Fluor channel (ΔF/A) and fitted with an exponential rise and decay (*74*). The APCaT peak ΔF/A amplitude and time integral values were derived from the fitted curves. The linearity of APCaT summation with increasing number of somatic APs was determined in each spine by normalizing the peak amplitude and time integral to the values of APCaTs elicited by single APs (1bAP). The distances from the soma to the parent dendritic segment of the imaged spines were determined *post hoc* using Simple Neurite Tracer (*75*) from three-dimensional reconstructions (Z-stacks) recorded in the AlexaFluor 594 channel at the end of the experiment. The spines located on the same segment were attributed the same distance from soma as the parent dendritic segment. The data were classified according to the somatic distance of their parent dendritic segment and grouped in three distance zones (in µm): 0 – 150, 150 – 250, and 250 – 400. The distribution of spine locations is shown in Fig. S5.

#### Immunohistochemistry

Rats were given a lethal dose of Euthatal, i.p. and transcardially perfused with iced-cold saline and 4% PFA in phosphate buffer (PB). Brains were removed and post-fixed in the same fixative overnight and were then immersed in 30% sucrose in PB for 48 hours. Free floating coronal sections were cut at 35µm thickness on a freezing microtome and stored in cryoprotective solution in −20°C freezer prior to immunohistochemistry. For total and phospho-ERK (active ERK) immunostaining, anti-p44/42 MAPK (#4695) and anti-phospho-p44/42 MAPK (#4370) primary antibody (1:1000 for both antibodies, Cell Signaling) were used, respectively. The detection of the bound antibodies was carried out using a standard peroxidase-based method (ABC-kit, Vectastain), followed by DAB (Sigma-Aldrich) and 3% H_2_O_2_ solution. Sections were then mounted on poly-L lysine coated slides, dehydrated, and mounted in DPX mounting medium (Sigma-Aldrich). For CREB and phosphorylated CREB immunofluorescence staining, anti-CREB and anti-phospho-CREB primary antibody (CREB (#9197): 1:1000; pCREB (#9198): 1:500; Cell Signaling) were used. Alexa Fluor 488 anti-rabbit secondary antibody (1:1000, Invitrogen, UK) was used to label the cells. DAPI (1:1000, Sigma-Aldrich) staining was also applied for nuclei staining. Sections were then mounted on poly-L lysine coated slides, dehydrated, and mounted with Mowiol mounting medium (Sigma-Aldrich). Images of the regions of interests were captured from both hemisphere from 3 to 4 sections (each 420µm apart) per animal. Coordinates of sections being processed was restricted to the dorsal hippocampus between bregma −2.76 mm and −3.96 mm (*76*). Images of DAB staining sections were captured using a 10x objective lens on Leica DMRB microscope with an Olympus DP70 camera. Images for immunofluorescence sections were captured using a 20x objective lens on Leica DM6000B upright time-lapse system. Data (i.e., positive cells counts per µm^2^) was quantified using Image/J-Fiji.

#### Intra-hippocampal infusion

Rats were surgically implanted with stainless steel double guide cannula (Plastics One, UK, 22 gauge, 3 mm below pedestal) under general anesthesia with Isoflurane (Isoflurane: 5% - 2%, flow rate: O_2_: 0.8 litter/min), as described previously (*77*). All surgery was carried out aseptically to minimize the risk of infection. Briefly, the double guide cannula was placed 3.8 mm apart and was aimed at the dorsal hippocampus 3.5mm posterior from the bregma. All rats were left to recover for a minimum of 6 days before behavioral experiments. Bilateral infusions via the chronically indwelling cannula were carried out in awake rats using a syringe pump, connected to injectors (28-gauge, projecting 1mm beyond the guide cannula) by polyethylene tubing. Diltiazem Hydrochloride (10 nmol/µl, DTZ, Sigma-Aldrich, UK) was dissolved in vehicle solution: 0.1M sterile phosphate-buffered saline (PBS), pH 7.2, treated with diethyl pyrocarbonate (DEPC). LM22B-10 (2µM, Tocris Bioscience, UK) was dissolved in vehicle PBS containing 0.1% DMSO. Each hemisphere received 1µl of drug solution or vehicle at the rate of 0.5µl per minute before the start of specific trials as indicated. The volume of LM22B-10 and PBS was 1µl per hemisphere and delivered at the rate of 0.5µl per minute. The injection cannula remained in place for further 2 mins to allow time for the diffusion of solutions away from the injection site before they were slowly removed. Histological determination of the cannula placement using β-thionin staining of Nissl substance showed the placement of cannula in the dorsal hippocampus with minimal tissue damage.

#### Systemic administration of drugs

LM22B-10 (Tocris Bioscience, UK) for peripheral injections was prepared as described (*55*). The animals were weighted and the amount of drug necessary to deliver 25 mg/kg in 1 ml injection volume was dissolved in HCl then mixed with 5% Cremophor (Sigma-Aldrich) and 2X sterile PBS. The solution was adjusted for pH 5 then diluted with double deionized water to 1 X PBS and 2.5% Cremophor. The drug solution was injected via intra-peritoneally 60 mins prior to each training stage of LI or before immunohistochemical and *ex vivo* slices preparations.

---

**Fig. S1.**
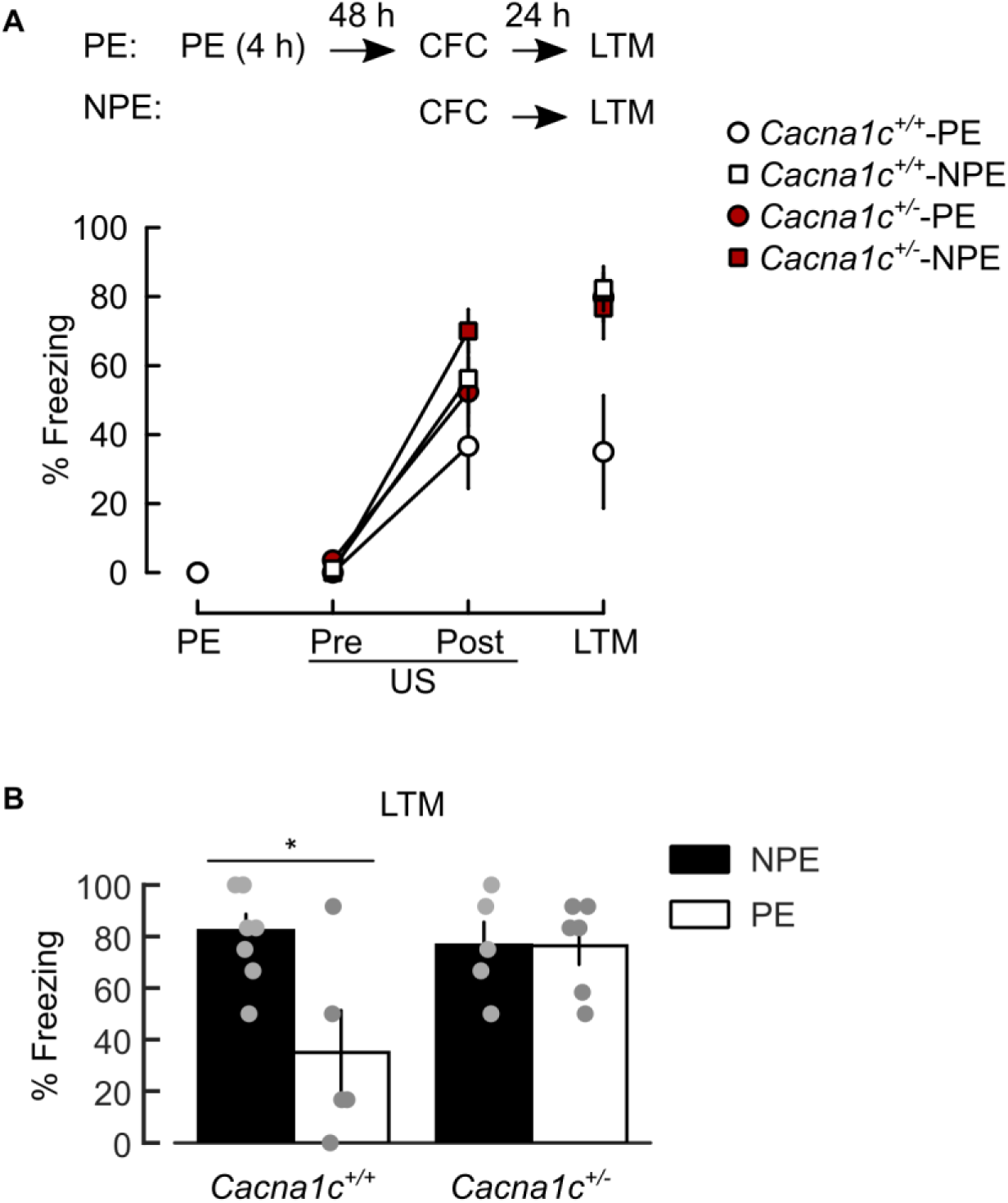
LI of CFM deficit in *Cacna1c^+/-^* animals replicated in a variation of the LI paradigm. (**A**) *Top:* schematics of the behavioral protocols; PE: 4h context pre-exposure (*Cacna1c^+/+^*-PE: n=5; *Cacna1c^+/-^*-PE: n=7); CFC: contextual fear conditioning comprising Pre-US and Post-US epochs before and after footshock (US), respectively; LTM: long-term memory test (CFM recall). NPE: non-pre-exposed (*Cacna1c^+/+^*-NPE: n=8; *Cacna1c^+/-^*-NPE: n=5). *Bottom*: Freezing responses at each session. The results showed a significant effect of trial (F_(2,42)_=74.671, p<0.0001), genotype (*Cacna1c^+/+^ versus Cacna1c^+/-^*: F_(1,21)_=6.521, p=0.018) and pre-exposure (PE *versus* NPE: F_(1,21)_=8.017, p=0.01), and a marginal interaction between group × pre-exposure interaction (F_(1,21)_=4.182, p=0.054). (**B**) Summary of freezing responses at LTM. There was significant effect of pre-exposure (F_(1,21)_=5.453, p=.003) and genotype × pre-exposure interaction (F_(1,21)_=7.088, p=.015) at LTM. Only in the *Cacna1c^+/+^* rats did PE reduce conditioned freezing at LTM1 indicating intact context LI. The high levels of CFM measured in *Cacna1c^+/-^* rats pre-exposed 48h before CFC was similar to that seen previously with a 24 h delay between PE and CFC (Figure 1). Data shown as means ±SEM. *p <0.05 determined by two-way repeated measures ANOVA followed by Tukey adjustment of p-values for contrasts.

**Fig. S2.**
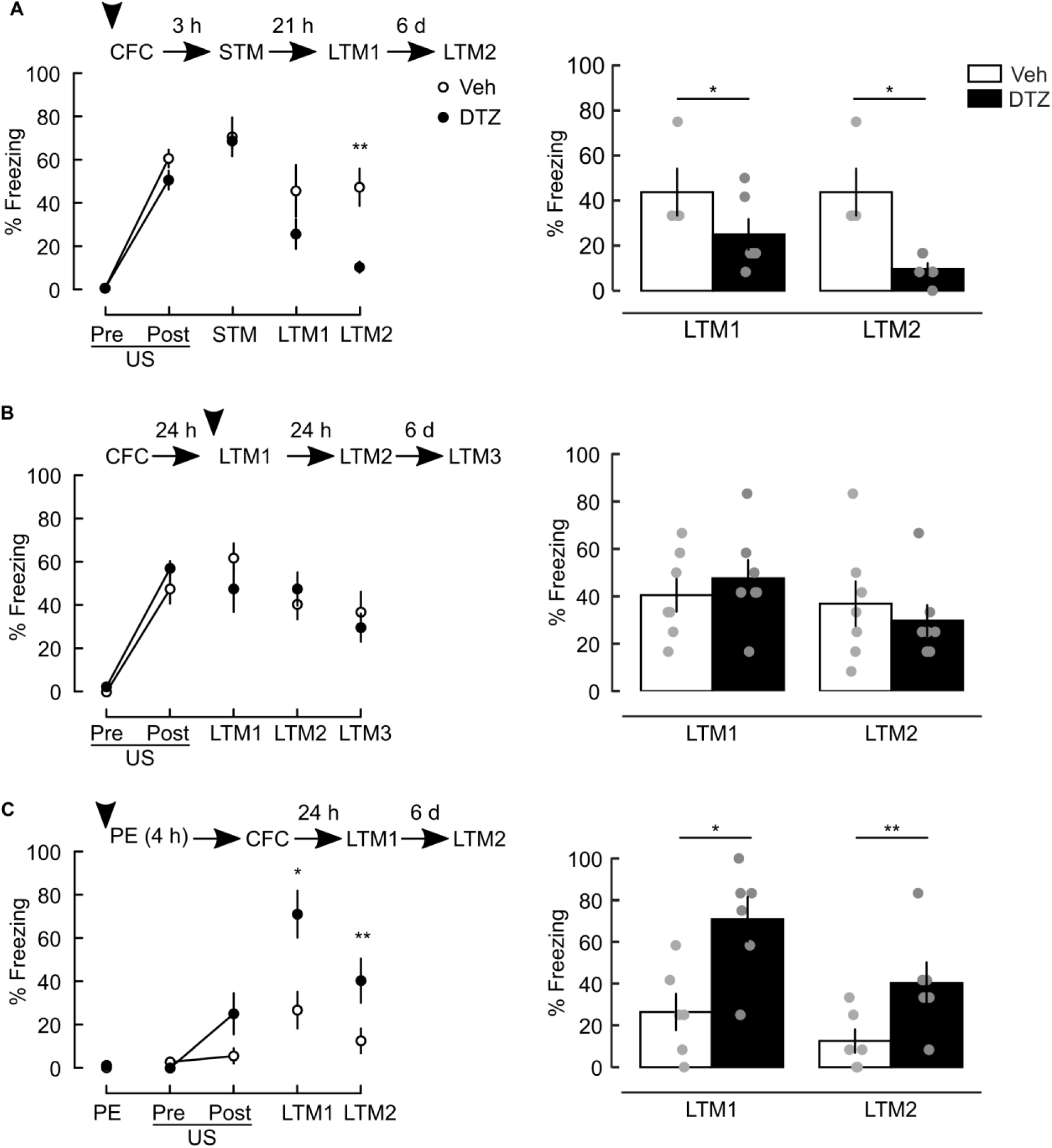
Intrahippocampal infusion of the L-type VGCC antagonist diltiazem disrupts components of contextual fear memory. **All panels:** *Left*: Plots of freezing response during the trials of the contextual fear conditioning paradigms, with session schematics at top. PE: 4h context pre-exposure; CFC: contextual fear conditioning comprising Pre-Us and Post-US epochs before and after footshock (US), respectively; STM, LTM1, LTM2, and LTM3: short-term and long-term contextual fear memory (CFM) retrieval tests. Arrowheads: bilateral infusion of 1µl diltiazem (DTZ, 10 nmol/µl) or vehicle (Veh) in dorsal hippocampus, 40 min before the indicated session. *Right*: Summary of the freezing responses at LTM tests. (**A**) DTZ infusion before CFC blocks the consolidation of CFM (treatment × trial: F_(4,36)_=4.371, p=0.006; pairwise comparison between treatments at LTM2: F_(4,36)_=8.246, p=0.002; DTZ: n=6, Veh: n=6). (**B**) There was no difference between the levels of freezing during CFC between the two groups (effects of treatment: F_(1,12)_=2.362, p=0.15; trial: F_(1,12)_=180.390, p<0.001; treatment × trial: F_(1,12)_=0.878, p=0.367). DTZ infusion prior to LTM1 does not affect retrieval of the CFM at LTM1 or in subsequent trials (effect of treatment F_(1,12)_=0.32, p=0.582, trial: F_(2,24)_=5.035, p=0.015; treatment × trial interaction: F_(2,24)_=1.305, p=0.290). DTZ: n=7; Veh: n=7. The effect of trial reflects the reduction in freezing response with multiple recalls indicative of normal between session extinction in the absence of DTZ. (**C**) DTZ infusion before pre-exposure blocks the establishment of latent inhibition of CFM (treatment × trial interaction: F_(4,40)_=5.381, p=0.001; treatment comparisons at LTM1: F_(4,40)_=10.364, p=0.009, LTM2: F_(4,40)_=5.9178, p=0.035; DTZ: n=6, Veh: n=6). Data shown as means ±SEM. *p <0.05, **p<0.01 determined by two-way repeated measures ANOVA followed by Tukey adjustment of p-values for contrasts.

**Fig. S3.**
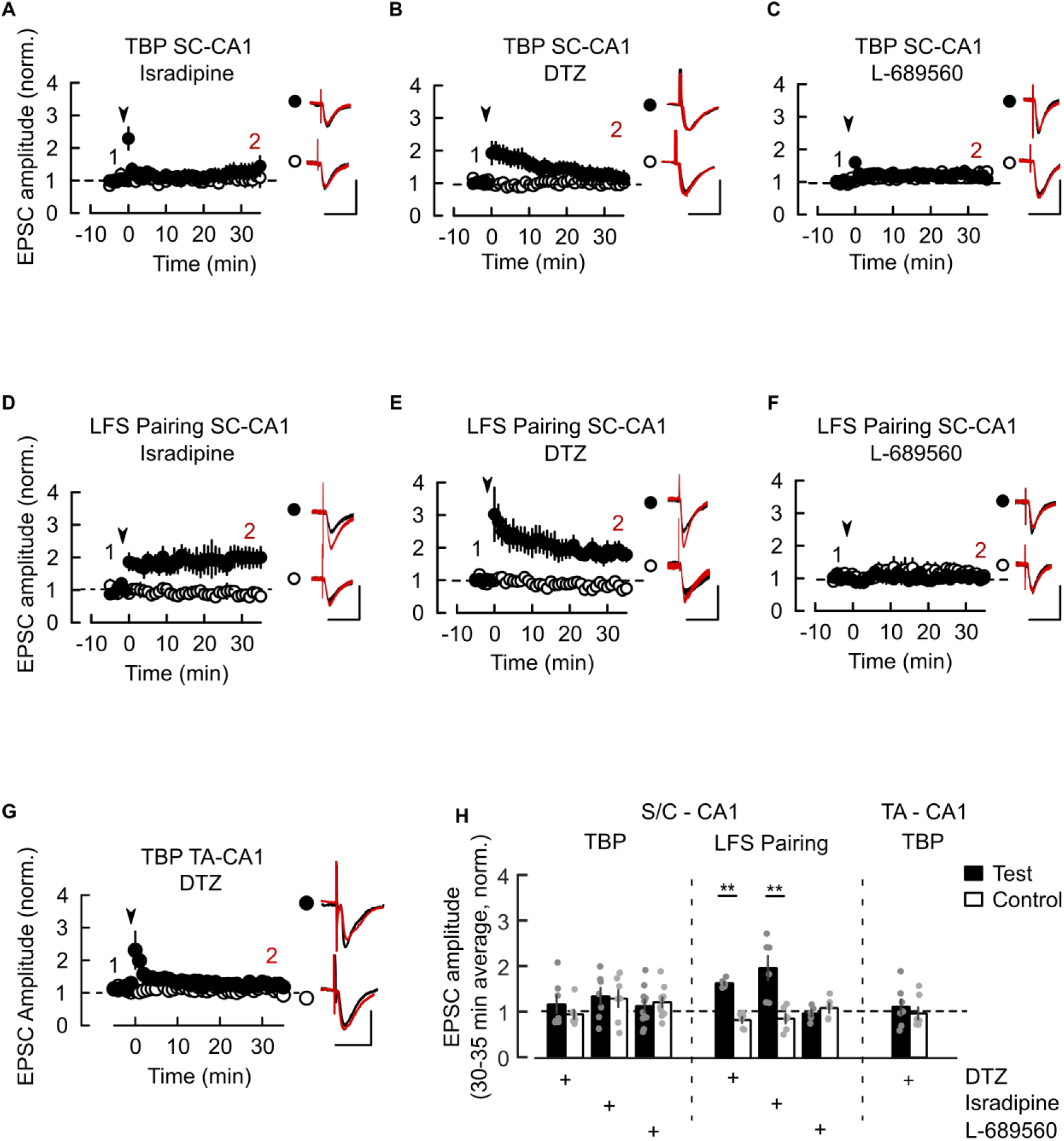
NMDAR and L-VGCC dependence of LTP at SC-CA1 and TA-CA1 synapses in *Cacna1c^+/+^ ex vivo* slices. (**A-C**) TBP-induced LTP at SC-CA1 synapses is blocked by L-VGCC antagonists Isradipine (10 µM, A) or diltiazem (DTZ, 100 µM, B) and by the NMDAR antagonist L-689560 (5 µM, C). (**D-F**) SC-CA1 LTP induced by LFS-pairing is insensitive to L-VGCC antagonists Isradipine (D) or DTZ (E) but is blocked by L-689560 (F). (**G**) TBP-induced LTP at TA-CA1 synapses is blocked by DTZ. Plots in A-G: Time course of EPSC amplitude normalized to 5 min before delivery of TBP protocol to the Test pathway (arrowheads). *Insets*: Average EPSC waveforms 5 min (1, black) and 30-35 min (2, red) after TBP. Scale bars: 50 pA, 50 ms. (**H**) Summary of normalized EPSC amplitude change 30-35 min after induction of LTP, in A-G. Pairwise Test vs Control for TBP-induced LTP at SC-CA1: Isradipine: p=0.99, n=7/5; DTZ: p=0.9, n=7/6; L-689560: p=0.927, n=10/6; LFS-pairing LTP in SC-CA1, Isradipine: p=0.0026, n=6/5; DTZ: p=0.0013, n=6/5; L-689560: p=0.9, n=5/4; TBP LTP at TA-CA1: DTZ: p=0.86, n=7/5. Sample sizes given as cells/animals. All drugs were bath applied throughout the recording. Data shown as means ±SEM. *p <0.05, **p<0.01 determined by two-way ordinal regression (cumulative link model) followed by analysis of deviance (ANODE).

**Fig. S4.**
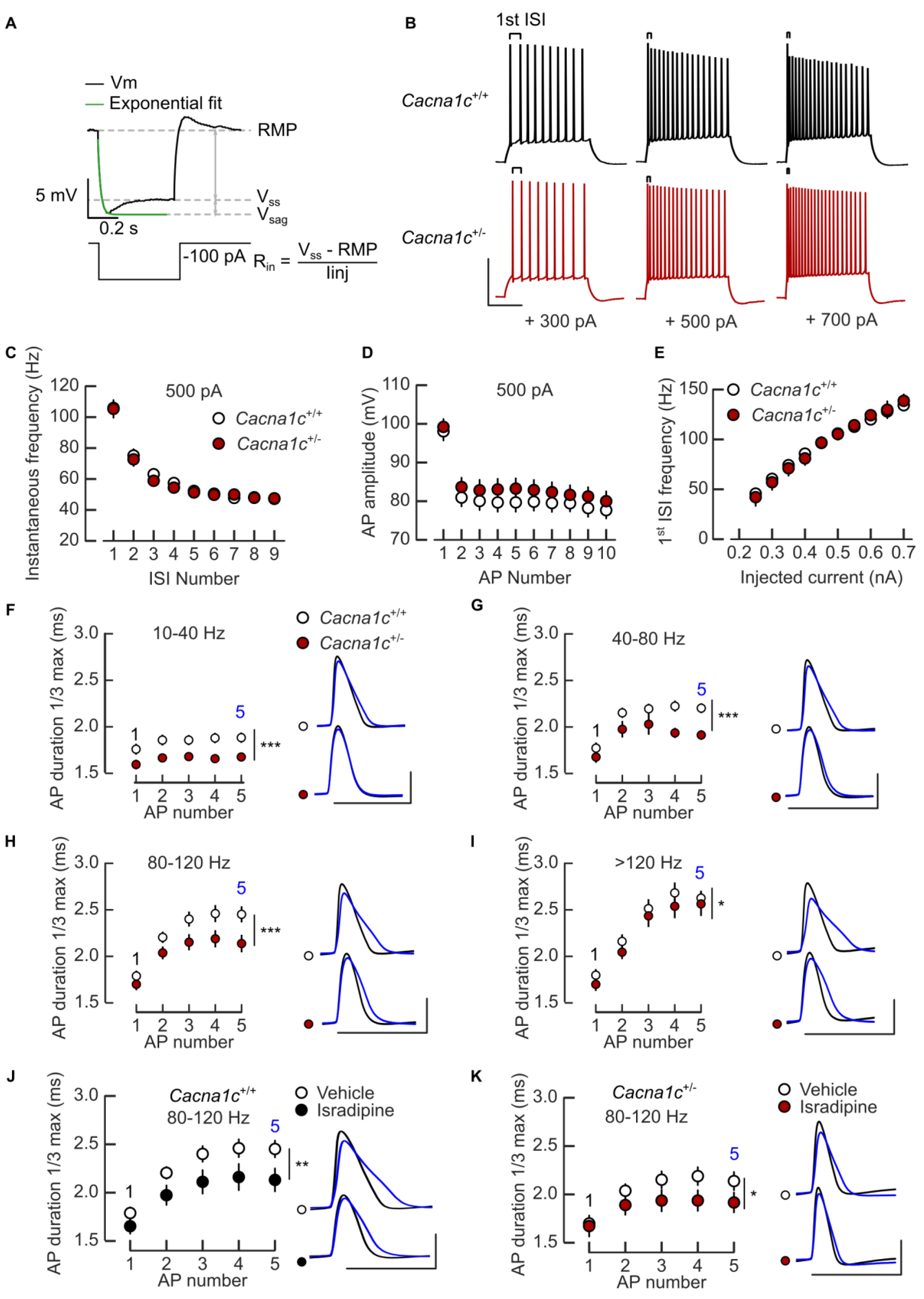
Reduced frequency-dependent broadening of somatic action potentials in CA1 pyramidal neurons from *Cacna1c^+/-^* animals. (**A**) Schematic of passive membrane properties measurement: resting membrane potential (RMP), input resistance (R_in_), V_sag_ and the exponential fit for membrane time constant (τ_m_) determination from membrane voltage recordings during hyperpolarizing current injections. (**B**) Examples of action potential (AP) trains discharged during 500 ms depolarising current (intensity shown underneath). ISI: inter-spike interval. (**C-E**) *Cacna1c^+/+^* and *Cacna1c^+/-^* neurons fire APs with comparable frequency adaptation (effect of spike number: LR_(9)_=328.27, p<0.0001; genotype: LR_(1)_=0.48, p=0.48, C), amplitude attenuation (effect of AP number: LR_(9)_=430.12, p<0.0001; genotype: LR_(1)_=0.11, p=0.73, D) and similar frequency – current intensity relationship (E). Data in C,D: first 10 AP fired during +500 pA current (500 ms); *Cacna1c^+/+^*: n=43/15; *Cacana1c^+/-^*: n=30/10 (cells/animals). (**F-I**) Frequency-dependent broadening of somatic APs is attenuated in *Cacna1c^+/-^ versus Cacna1c^+/+^* neurons. Genotype effect at 10-40 Hz (F): LR_(1)_=30.67 p<0.0001; 40-80 Hz (G): LR_(1)_=41.12, p< 0.0001; 80-120 Hz (H): LR_(1)_=22.83, p<0.0001; >120 Hz (I): LR_(1)_ = 5.8, p=0.015. Sample sizes per frequency band respectively, for *Cacna1c^+/+^* and *Canca1c^+/-^*, at 10-40 Hz: n=36/15 and n=26/10; 40-80 Hz: n=30/15 and n=29/10; 80-120 Hz: n=43/15 and n=28/10; >120 Hz: n=37/15 and n=26/10. (**J,K**) Isradipine (10 µM) inhibits AP broadening at 80-120 Hz in both *Cacna1c^+/+^* (LR_(1)_ = 7.16, p=0.007, n=26/11, J) and *Cacna1c^+/-^* (LR_(1)_=5.63, p=0.017, n=13/3, K) neurons. Plots in F-K: AP durations for the first 5 AP. Insets: 1^st^ (2, black) and 5^th^ (5, blue) AP waveform; scale bars: 5 ms, 50 mV. When used, Isradipine was bath applied throughout the experiment. Data shown as means ±SEM. *p <0.05, **p < 0.01 and ***p< 0.001, determined by two-way repeated measures ANOVA followed by Tukey adjustment of p-values for contrasts.

**Fig. S5.**
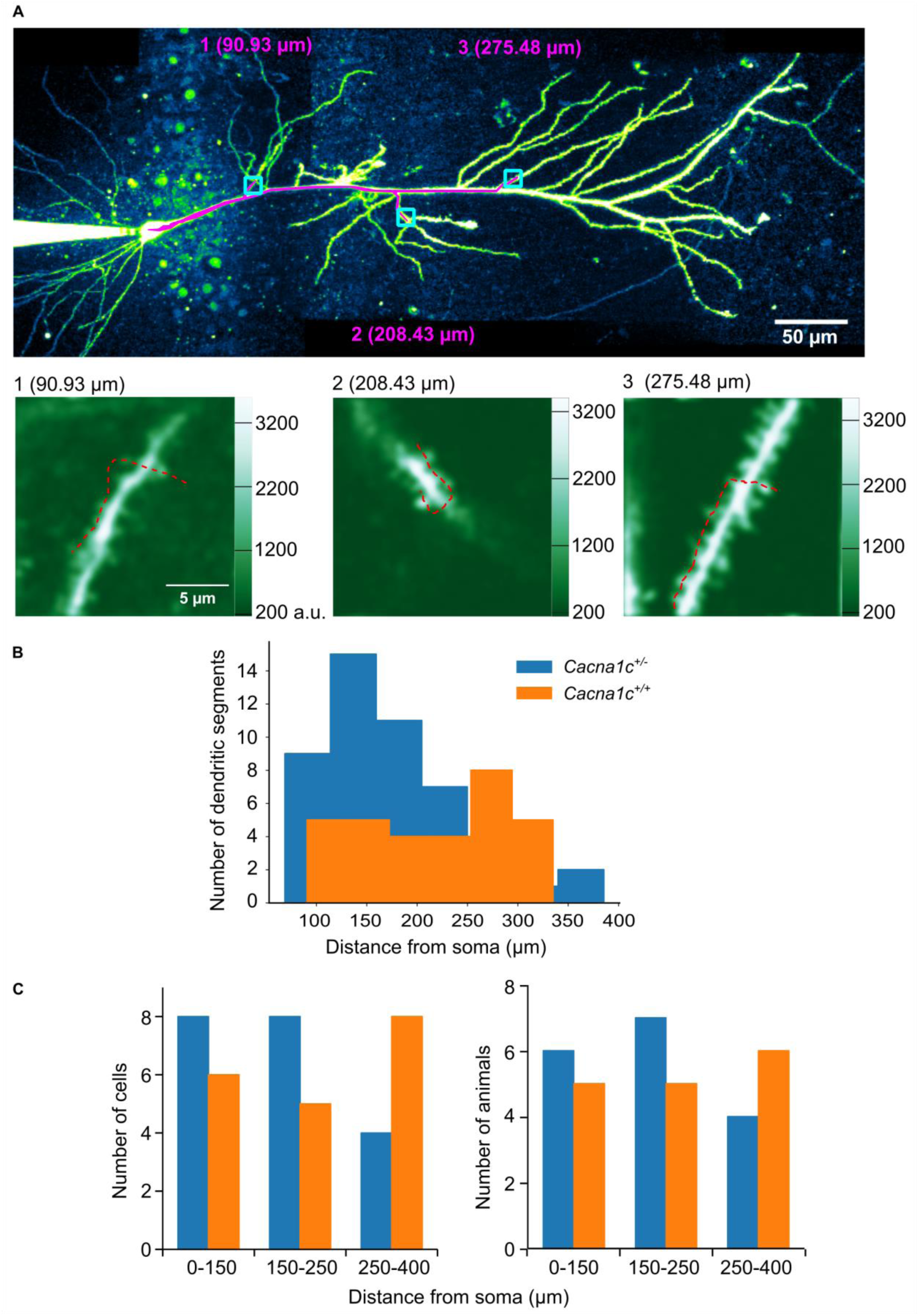
Location of imaged spines. (**A**) *Top*: Z-projected image of a CA1 pyramidal neuron (AlexaFluor 594 channel) with the overlaid “paths” (pink) used to measure the distance from the soma to three imaging locations (blue squares). The “paths” were traced and measured using the Single Neurite Tracer (Image/J). *Bottom*: Raster scans through the imaging locations shown in the Z-projected image (trajectory of line scan overlaid in red); a.u.: arbitrary units. Imaging was performed on spines on a single dendritic segment at a given location. Imaging locations were selected pseudo-randomly, ensuring that the imaged spines were located on distinct dendritic branches originating from the main trunk at least 30 µm apart. (**B**) Distribution of soma-spine distances. Spines on the same dendritic segment were attributed the same distance from the soma. (**C**) Distribution of the somatic distance zones across cells (left) and animals (right) in both genotypes.

**Figure S6.**
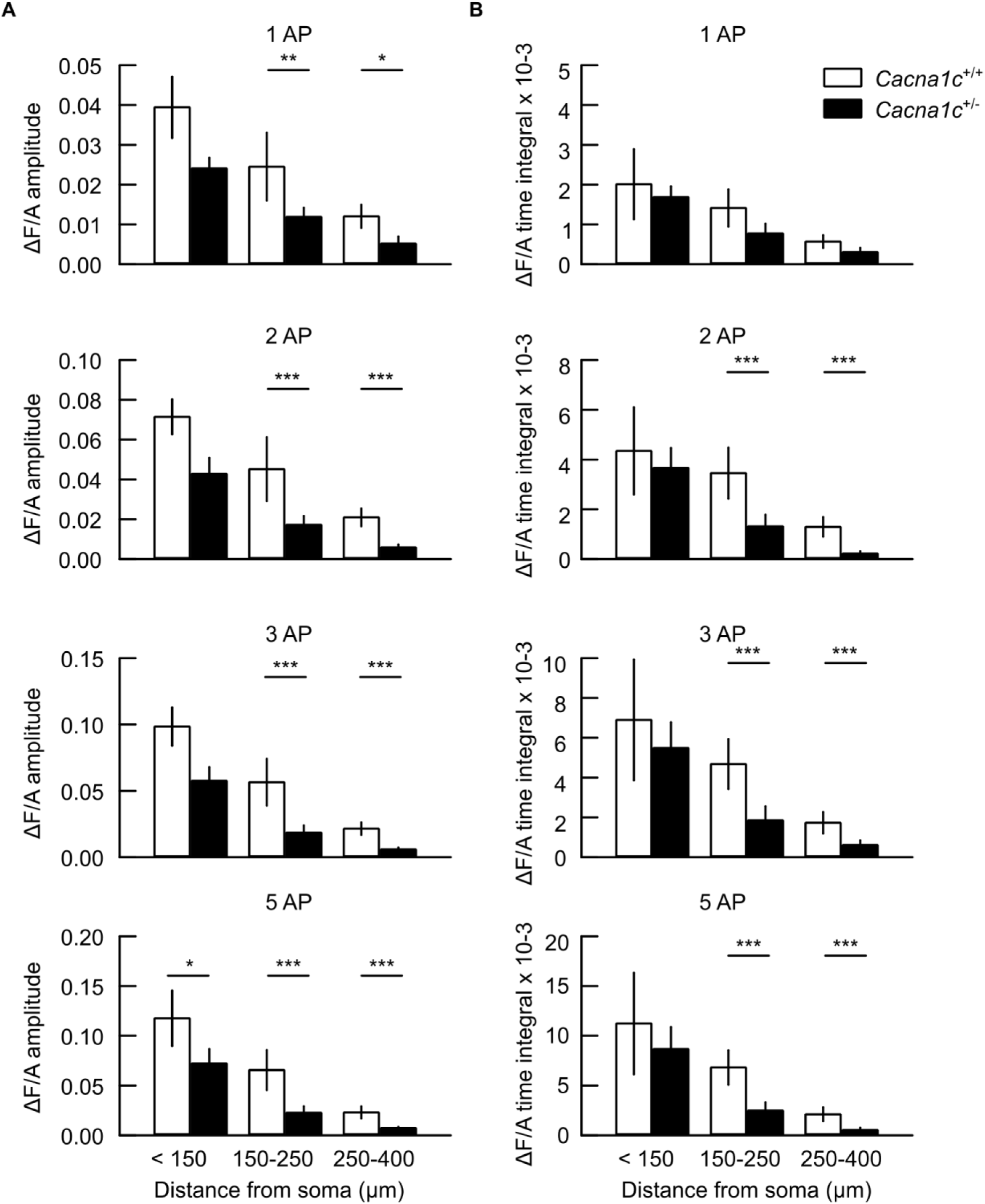
Attenuation of spine calcium transients in CA1 pyramidal cells with distance from soma. (**A,B**) Summary of peak ΔF/A amplitude (A) or ΔF/A time integral (B) of spine Ca^2+^ transients (APCaTs) evoked by 1, 2, 3 or 5 back-propagated action potentials (AP, 100 Hz intra-burst, top to bottom) *versus* spine distance from soma. APCaTs evoked with bursts of two to five APs are significantly smaller in spines located beyond 150 µm from soma, in *Cacna1c^+/-^* compared to *Cacna1c^+/+^* neurons. Statistical analysis results are given in Supplementary Table S2; all p-values for pairwise comparisons are given in Supplementary Table S3. Data for the 5 AP APCaTs are reproduced from Fig. 3D,E. Data shown as means ±SEM. *p <0.05 and **p < 0.01, determined by two-way repeated measures ANOVA followed by Tukey adjustment of p-values for contrasts.

**Figure S7.**
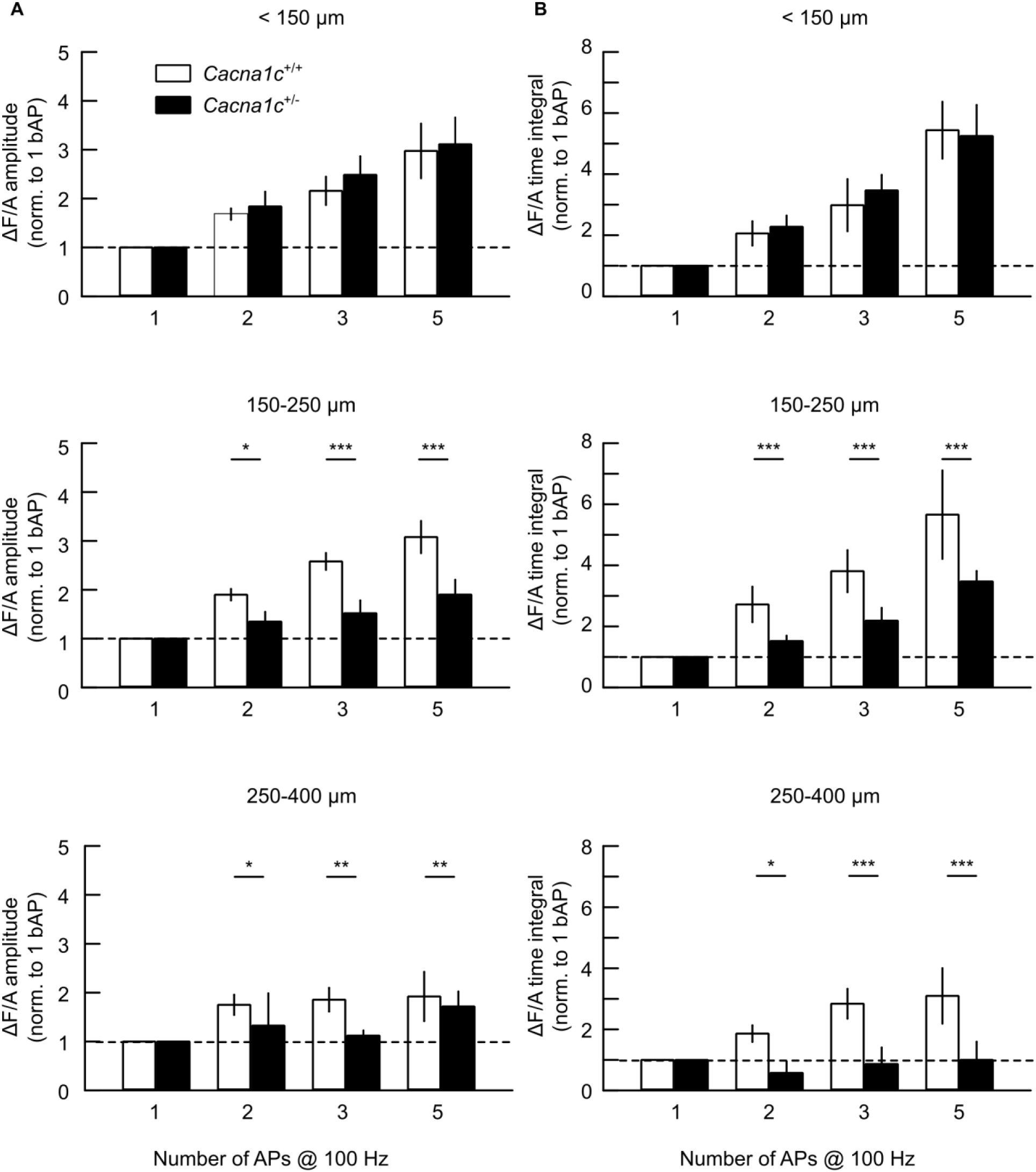
Summation of back action potential-mediated calcium transients (APCaTs) in pyramidal CA1 neurons. (**A,B**) Summary of peak ΔF/A amplitude (A) or ΔF/A time integral (B) of spine Ca^2+^ transients (APCaTs) evoked by 1, 2, 3 or 5 back-propagated action potentials (AP, 100 Hz intra-burst) normalized to the values at 1 AP, in the three distance zones analyzed (top-to-bottom). In spines located beyond 150 µm from the soma, the summation of APCaTs with the number of APs is reduced in *Cacna1c^+/-^ versus Cacna1c^+/+^* neurons. Statistical analysis results are given in Supplementary Table S2; all p-values for pairwise comparisons are given in Supplementary Table S3. Middle panels (150-250 µm) are reproduced from Fig. 3F,G. Data shown as means ±SEM. *p <0.05, **p < 0.01 and ***p<0.001 determined by two-way repeated measures ANOVA followed by Tukey adjustment of p-values for contrasts.

**Figure S8.**
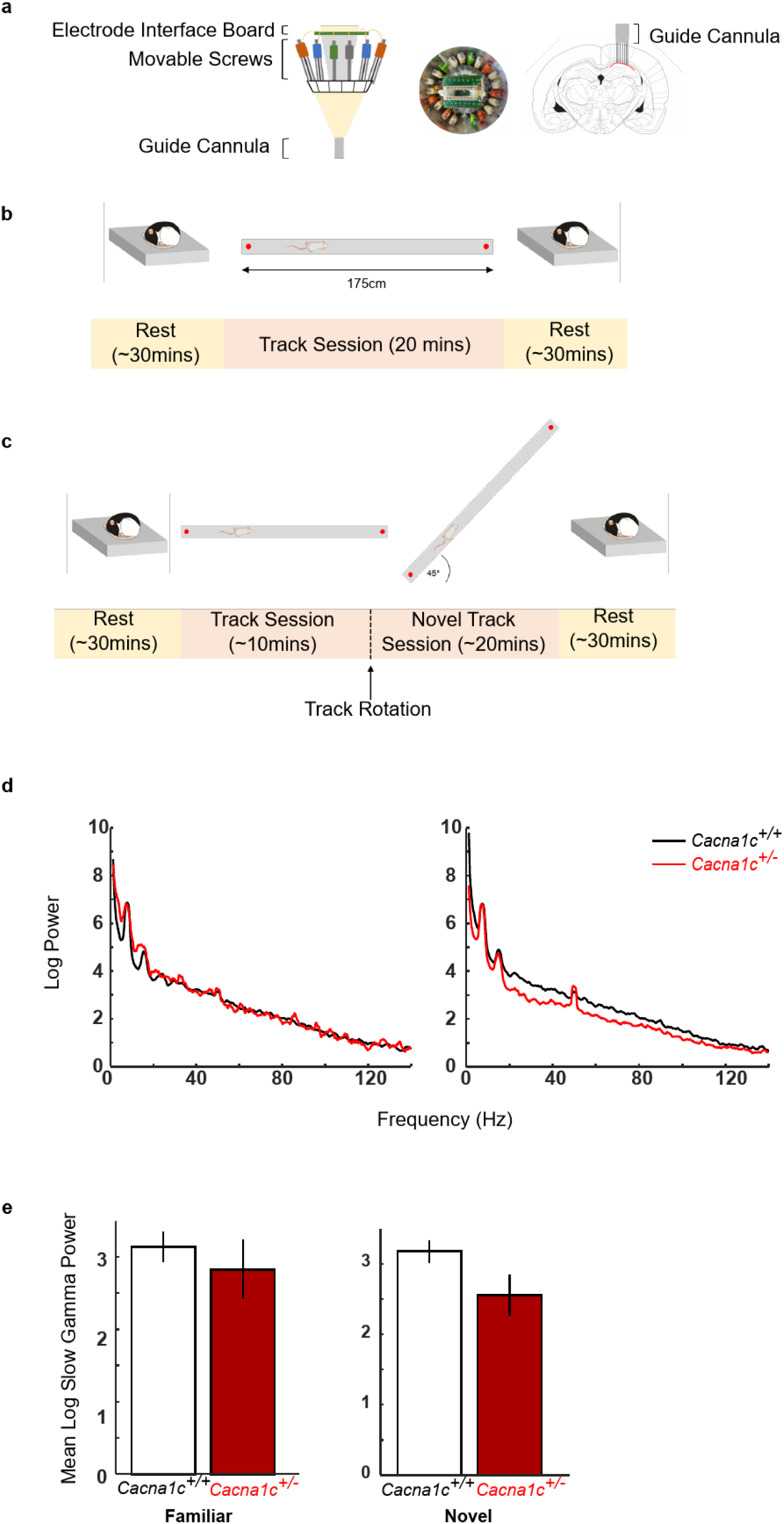
Local field potential recordings in dorsal CA1 during novel vs familiar track behavioral protocol. (**A**) Tetrode Microdrive used for recording in vivo local field potentials. Left: Schematic of Microdrive viewed from the side. Middle: Photo of Microdrive viewed from the top. Right: Position of guide cannula during surgery and targeting of tetrodes to dorsal CA1 of hippocampus. Red line indicates dorsal CA1 cell layer. (**B**) Familiar track behavioral protocol. Following a ∼30-minute rest period, rats freely run back and forth on the linear track for sucrose rewards followed by another ∼30-minute rest period. (**C**) Novel Track Behavioral Protocol. As (B) but midway through the track session, rats were taken off the track and the track was rotated 45° before being placed back on to continue running freely for sucrose rewards. (**D**) Power spectra taken from track runs on the familiar (left) and novel track (right). (**E**) Mean slow gamma (25-55Hz) power did not differ on the familiar (p = 0.5607; Student’s t-test) or novel track (p = 0.1306; Student’s t-test). (*Cacna1c^+/+^*: n = 5; *Cacna1c^+/-^*: n=7; Data presented as mean ± SEM).

**Figure S9.**
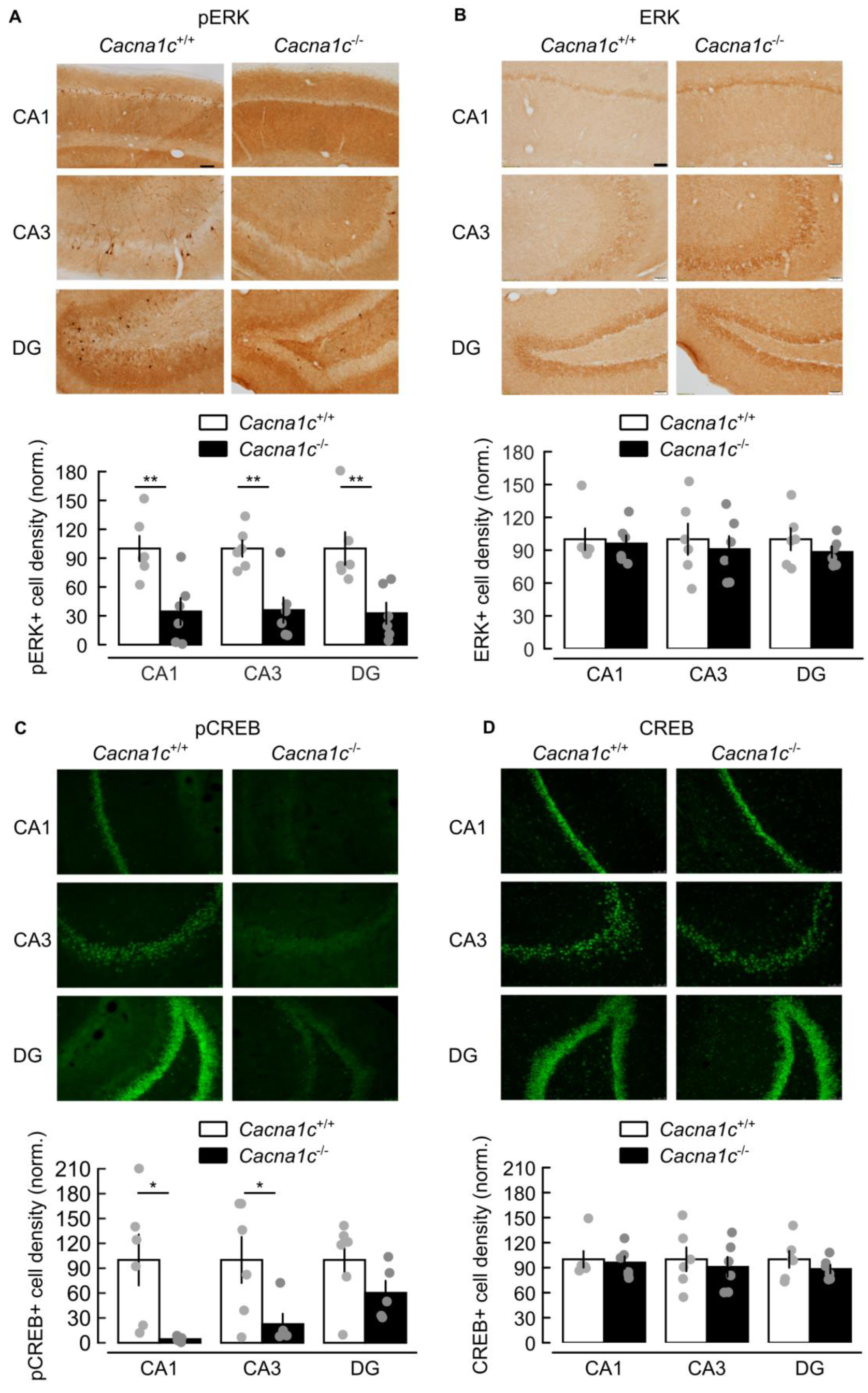
Reduced basal levels of phosphorylated ERK and phosphorylated CREB in the hippocampus of *Cacna1c^+/-^* rats. (**A**) Baseline levels of pERK are reduced in the hippocampus of *Cacna1c^+/-^* compared to *Cacna1c^+/+^* rats. Effects of genotype: F_(1,10)_=14.617, p=0.003 and region: F_(2,20)_=0.038, p=0.962; genotype × region interaction: F_(2,20)_=0.038, p=0.962; independent sample t-tests for pERK-positive cell density in CA1: t_(10)_=3.423, p=0.007; CA3: t_(10)_ = 4.095, p=0.002; DG: t_(10)_=3.311, p=0.008. (**B**) Similar levels of total ERK expression in both genotypes (F < 1). (**C**) Reduced baseline hippocampal levels of pCREB in *Cacna1c^+/-^* compared to *Cacna1c^+/+^* rats. Effects of genotype: F_(1,9)_=6.719, p=0.029 and region: F_(2,18)_=2.524, p=0.108; genotype × region: F_(2,18)_=2.524, p=0.108; independent sample t-tests for pCREB-positive cell density in CA1: t_(5.019)_=3.105, p=0.027; CA3: t(6.875) = 2.537, p = 0.039; DG: t_(10)_=1.549, p=0.156. (**D**) Levels of total CREB expression are similar in both genotypes (F < 1). **All panels**: *Top*: examples of immunoperoxidase (A,B) and immunofluorescence (C,D) staining; *Bottom*: summary of immunopositive cell densities (mean ± SEM), in the dorsal hippocampal regions CA1, CA2 and dentate gyrus (DG), normalized to the average value in *Cacna1c^+/+^* tissue. Sample sizes: *Cacna1c^+/+^* (n = 6) and *Cacna1c^+/-^* (n=6) animals. Scale bars: immunoperoxidase images: 100 µm; immunofluorescence: 50 µm. Data shown as means ±SEM. *p <0.05 and **p < 0.01, two-way repeated measures ANOVA followed by independent sample t-test.

**Figure S10.**
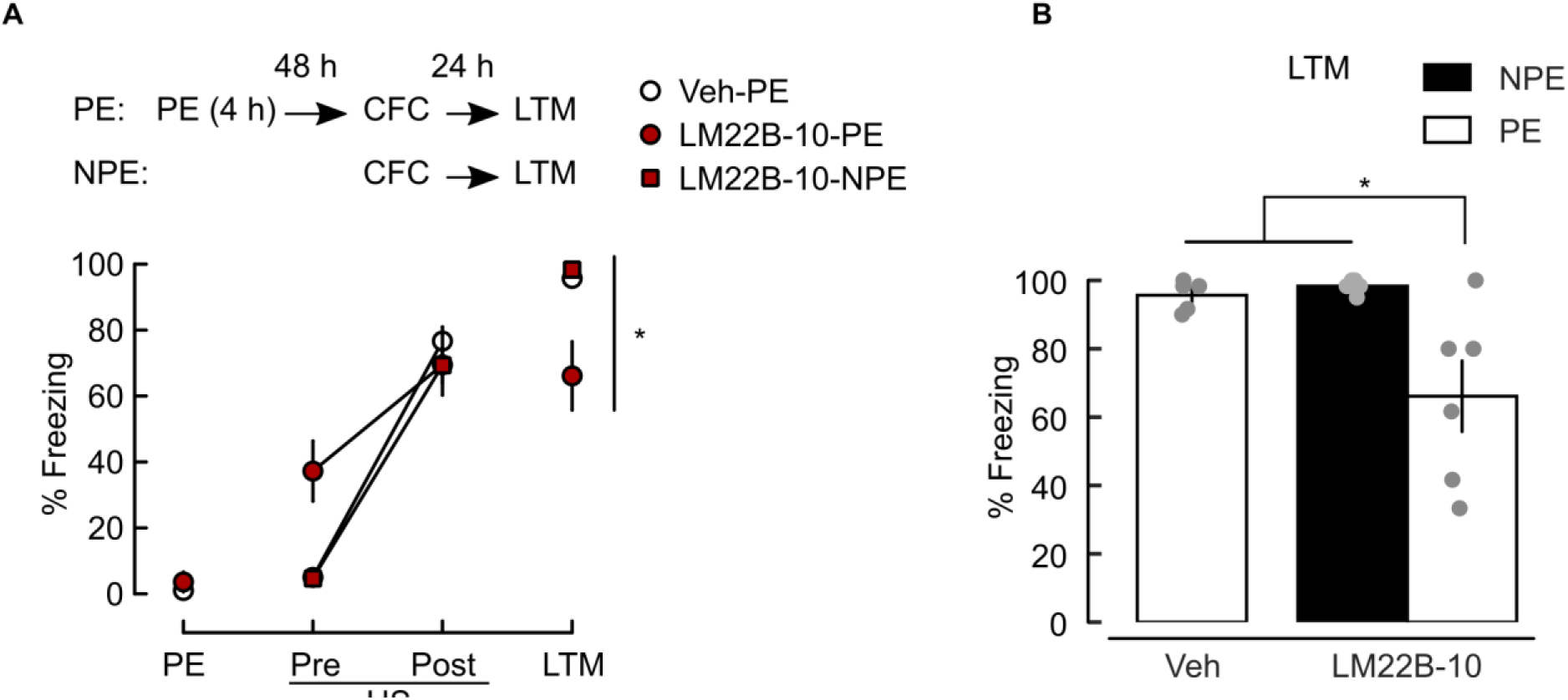
Systemic administration of LM22B-10 rescues LI of CFM in *Cacna1c^+/-^* animals. (**A**)Top schematic: paradigm sessions, with notations as in Fig. S1. Animals received an i.p. bolus of either LM22B-10 (25 mg/kg; pre-exposed: LM22B-10-PE, n=6; non pre-exposed: LM22B-10-NPE, n=6) or vehicle (pre-exposed: Veh-PE, n=5) 60 min before each stage. This schedule was chosen to avoid effects of state-dependency. Significantly different freezing behavior in LM22B-10-PE *versus* Veh-PE animals (treatment × trial interaction: F_(3,24)_=11.653, p<.0001). Freezing behavior at Pre-US trial is stronger in LM22B-10-PE animals *versus* Veh-PE (F_(3,24)_=7.858, p=0.023). (**B**) Freezing response at LTM is significantly smaller in LM22B-10-PE *versus* Veh-PE animals (F_(3,24)_=5.638, p=0.045) and *versus* LM22B-10-NPE animals (t_(5.0582)_=3.099, p=0.027). Data shown as means ±SEM. *p <0.05, determined by univariate ANOVA and independent sample t-test.

**Table S1.** Passive membrane and action potential firing properties of CA1 pyramidal neurons in *Cacna1c*^+/+^ and *Cacna1c*^+/-^ littermates.

| **Passive membrane properties:** | *Cacna1c*<sup>+/+</sup> | *Cacna1c*<sup>+/-</sup> | *p*<sup>†</sup> (U) |
| --- | --- | --- | --- |
| RMP (mV) | -71.77 ± 0.35 | -70.99 ± 0.45 | 0.39 (310) |
| *R*<sub>in</sub> (MΩ) | 66.57 ± 3.46 | 62.0 ± 3.22 | 0.6 (345) |
| *τ*<sub>M</sub> (ms) | 20.3 ± 1.3 | 23 ± 1.5 | 0.15 (337) |
| % Sag | 2.05 ± 0.17 | 1.94 ± 0.11 | 0.85 (261) |
| % Rebound | 10.35 ± 0.52 | 9.82 ± 0.39 | 0.6 (245) |
| **Action potential (AP) firing properties:** |  |  |  |
| *I*<sub>rh</sub> (pA) | 201.55 ± 11.36 | 237.55 ± 17.13 | 0.12 (511) |
| Number of AP<sup>‡</sup> | 18.99 ± 0.79 | 16.72 ± 1.22 | 0.23 (751) |
| Firing threshold (mV)<sup>‡</sup> | -54 ± 1.07 | -54.55 ± 0.77 | 0.57 (595) |
| First AP amplitude (mV)<sup>‡</sup> | 99.99 ± 1.48 | 99.22 ± 1.31 | 0.42 (693) |
| First AP maximum dV/dt (V/s)<sup>‡</sup> | 390.31 ± 14.24 | 400.27 ± 14.01 | 0.54 (591) |

† *p*-value and U statistic, two-sided Mann-Whitney U test, *Cacna1c*<sup>+/+</sup>: n=43/15; *Cacna1c*<sup>+/-</sup>: n=30/10.
‡ measured at 500 pA current injection.

Table S2: Statistical comparisons of APCaTs by regions of distance from soma. File Tigaret et al Table S2.xlsx

Table S3: P-values for pairwise comparisons of APCaTs in Cacna1c^+/+^ vs Cacna1c^+/-^ CA1 pyramidal neurons. File Tigaret et al table S3.xlsx

